# A massively parallel reporter screen defines the syntax of SECReTE, a secretion-enhancing *cis* regulatory targeting element, in mammalian cells

**DOI:** 10.64898/2026.10.12.758471

**Authors:** Gal Haimovich, Gabriela Lobinsky, Shu-Ting You, Dvir Schirman, Frida Grynspan, Ariel Gilert, Orna Dahan, Tsviya Olender, Jeffrey E. Gerst, Yitzhak Pilpel

## Abstract

Messenger RNA (mRNA) localization to specific subcellular regions is crucial for proper protein translation and localization. Potential means to localize mRNAs include *cis*-acting or structural motifs embedded within the sequence. We previously identified a <u>s</u>ecretion-<u>e</u>nhancing *c<u>is</u>* <u>r</u>egulatory <u>t</u>argeting <u>e</u>lement (SECReTE) enriched in mRNAs encoding secreted proteins from prokaryotes, eukaryotes, and viruses, found to enhance both protein synthesis and secretion by localizing mRNAs to the endoplasmic reticulum (ER) membrane. Defined as at least 10 consecutive *NNY* triplets (where *N* is any nucleotide and *Y* is a pyrimidine), this highly degenerate motif can be specified by approximately 10^15^ different sequence combinations. Yet, some combinations may prove more efficient than others in regulating mRNA localization and protein secretion.

To better understand SECReTE syntax and to identify robust combinations, we designed and synthesized a library of ∼5 × 10^4^ variants and inserted them into the 3’ untranslated region (3’UTR) of two reporter genes, green fluorescent protein (GFP) and immunoglobulin light chain. We expressed each variant in Chinese hamster ovary cells and assayed their activity using parallel assays to measure the effect of each variant upon mRNA partitioning between the ER and cytoplasm, and protein secretion. Our results reveal that ER-associated mRNAs are enriched with SECReTE elements of a defined syntax. Notably, *RRY* repeats (where *R* is purine) were superior to *NNY* repeats. Variants conferring a high mRNA ER localization score consistently enhanced protein secretion, confirming the mechanistic relationship between mRNA targeting to the ER and secretion efficiency. The correlation between variant performances across both reporters suggests a shared regulatory mechanism. Additional analysis identified a novel secretion-promoting *cis* element, called “split NNS” (where S is G or C). This work refines the syntactic definition of mRNA localization motifs that regulate protein secretion in both natural and heterologous expression systems.

## Introduction

The targeting of proteins to their correct intracellular or extracellular locations depends in many cases upon specific amino acid sequence motifs (*i.e.* protein targeting signals) embedded within the protein as well as mRNA *cis* elements that are required for local sub-cellular translation. In eukaryotes, proteins targeted to the secretory pathway (herein referred to as the secretome) are translated on the membrane of the endoplasmic reticulum (ER). Some secreted proteins are targeted to the ER via the signal peptide [SP] (1) located at the N-terminus of the protein. Once translated, this peptide is recognized by the signal recognition particle (SRP) on the ER membrane, which historically was thought to be responsible to tether the translating mRNA to the ER (2). Nuclear proteins are imported to the nucleus via a nuclear localization signal (NLS) (3). In addition, an N-terminal mitochondrial targeting signal [MTS] (4) peptide found in many nuclear-encoded mitochondrial proteins is recognized by the translocase of the outer membrane (TOM) complex, which then tethers the translating mRNA-ribosome complex to the mitochondria (5), while specific import complex proteins are thought to facilitate the import of peroxisomal targeting signal [PTS] -containing proteins into peroxisomes (6).

In addition to peptide sequences, the localization of mRNAs to specific sub-cellular compartments or organelles has been demonstrated in a wide range of cellular and organismal systems (7–10). This may allow for localized translation and, thereby, either facilitate or circumvent the need for direct protein targeting (9). In many cases, the mRNA localization is determined by specific and unique *cis* elements, generally termed ‘ZIP codes’ (8). For example, *ASH1* mRNA in budding yeast contains ZIP codes that target it to the bud tip (11), β-actin mRNA in animal cells contains ZIP codes that target it to the leading edge of fibroblasts (12), glutelin and prolamine mRNAs in rice endosperm cells contain ZIP codes that target them to specific sub-domains of the ER (13), and many neuronal mRNAs contain ZIP codes in their 3’ untranslated region (3’UTR) that control transport from the cell soma to neurites (14). RNA localization is also facilitated by RNA-binding proteins (RBPs) that recognize ZIP codes and/or structural elements within the mRNA and help target the RNA to the organelle. For example, some nuclear-encoded mitochondrial mRNAs are localized to the mitochondria by association of RBPs (*e.g.* Puf3p in yeast, CLUH in mice and PUM2 in human) that recognize specific sequences in their 3’UTR with the outer mitochondrial membrane (5). In addition, numerous ER-associated RBPs have been identified in yeast and mammals (*e.g.* P180/RRBP1, HDLBP/Vigilin, TIS11B and She2p) and their mutation can lead to mRNA mistrafficking and reduced protein expression and secretion (15–18). Despite these advances, RNA localization to organelles is not fully understood in terms of all the RBPs that may participate therein as well as the sequence and structural motifs present in the mRNAs which may not be readily discernable.

Work from our labs and others’ suggested that secretome-encoding mRNAs in yeast and mammals are found on the ER membrane prior to translation initiation, upon translation inhibition, or in the absence of a functional SRP (19–25). In our studies on RNA localization to the ER, we identified a <u>s</u>ecretion-<u>e</u>nhancing *<u>c</u>is* <u>r</u>egulatory <u>t</u>argeting <u>e</u>lement (SECReTE), which is enriched in secretome mRNAs in all organisms from prokaryotes to eukaryotes. In yeast, presence of the SECReTE motif was shown to enhance the synthesis and secretion of secreted proteins, such as Suc2 and Hsp150 (19). In contrast, mutations that eliminated the SECReTE motif reduced protein synthesis and secretion, and affected cell physiology (19). The motif consists of triplets of nucleotides in the form of *NNY* or *NYN*, where *N* is any nucleotide, and *Y* is a pyrimidine (*T* or *C*). Based on our SECReTE enrichment analysis in yeast, we set the threshold for a minimal SECReTE at 10 consecutive repeats of the (in-frame) triplet *NNY* or 10 consecutive repeats on the in-frame triplet *NYN*. We note though that the motif can be located both within the coding and non-coding (*e.g.* UTRs) regions of an mRNA. Interestingly hydrophobic amino acids are encoded by either U or C in the second codon position, hence the SECReTE motif of the in-frame *NYN* form is enriched within the coding sequences of the SP and transmembrane domains (TMD). In contrast, as secreted proteins do not tend to have consecutive runs of hydrophobic amino acids, they tend to have the second version of the SECReTE motif, the in-frame *NNY* from, which places the requirement for a pyrimidine nucleotide at the wobble position of the codons. Irrespective of the reading frame, the two versions of SECReTE syntax (*i.e. NYN* and *NNY*) are identical as they both feature pyrimidines spaced 3 nucleotides apart and embedded within an otherwise unspecified sequence.

Given the large number of consecutive triplet nucleotide repeats in the defined SECReTE motif, there is an exponential number of actual sequences that can confer it - for example, 10 repetitions of the *NNY* triplet yields 32^10^ = >10^15^ possible sequences. We identified ∼50,000 SECReTE variants present within the human protein coding genes (19), with about half present in secretome mRNAs (**File S1**). A more precise understanding of SECReTE syntax might help narrow down the key elements which control the production and secretion of a given secreted protein. This will be inherently useful for the heterologous production of proteins (*e.g.* hormones, growth factors, antibodies, *etc*.) for pharma and food industry applications.

To better define the syntax of SECReTE, we developed and tested two complementary high-throughput methods to scan and score tens of thousands of designed and natural variants (**Fig 1**) for mRNA ER localization and protein secretion. Briefly, we designed a synthetic DNA library of 47,975 SECReTE and non-SECReTE variants, that include both natural sequences, designed sequences, and sequences that tile secreted proteins that have or do not have a SECReTE motif. Each of these designed sequences was inserted at the 3’UTR of two reporter genes: one encoding a cell surface-expressed GFP and the other a cell surface-expressed, FLAG-tagged, IgG light chain (LC). The LC construct was meant to serve as a proxy for a real-world challenge of IgG bioproduction. Chinese hamster ovary (CHO) cells were transduced separately with either of these two genes libraries using a lentiviral system, such that every transduced cell gets one library variant. We scored each library variant in two massively parallel screening assays. An assay for reporter protein secretion conferred by each variant was based on quantification of surface protein levels. A second assay, based on cell fractionation, measured mRNA enrichment in the membrane fraction *vs*. cytoplasmic fraction of the cells. By comparing the enrichment of thousands of variants, we found several possible sequence-based syntax rules for successful SECReTE motifs that confer better mRNA ER localization and/or protein secretion.

**Fig 1.**
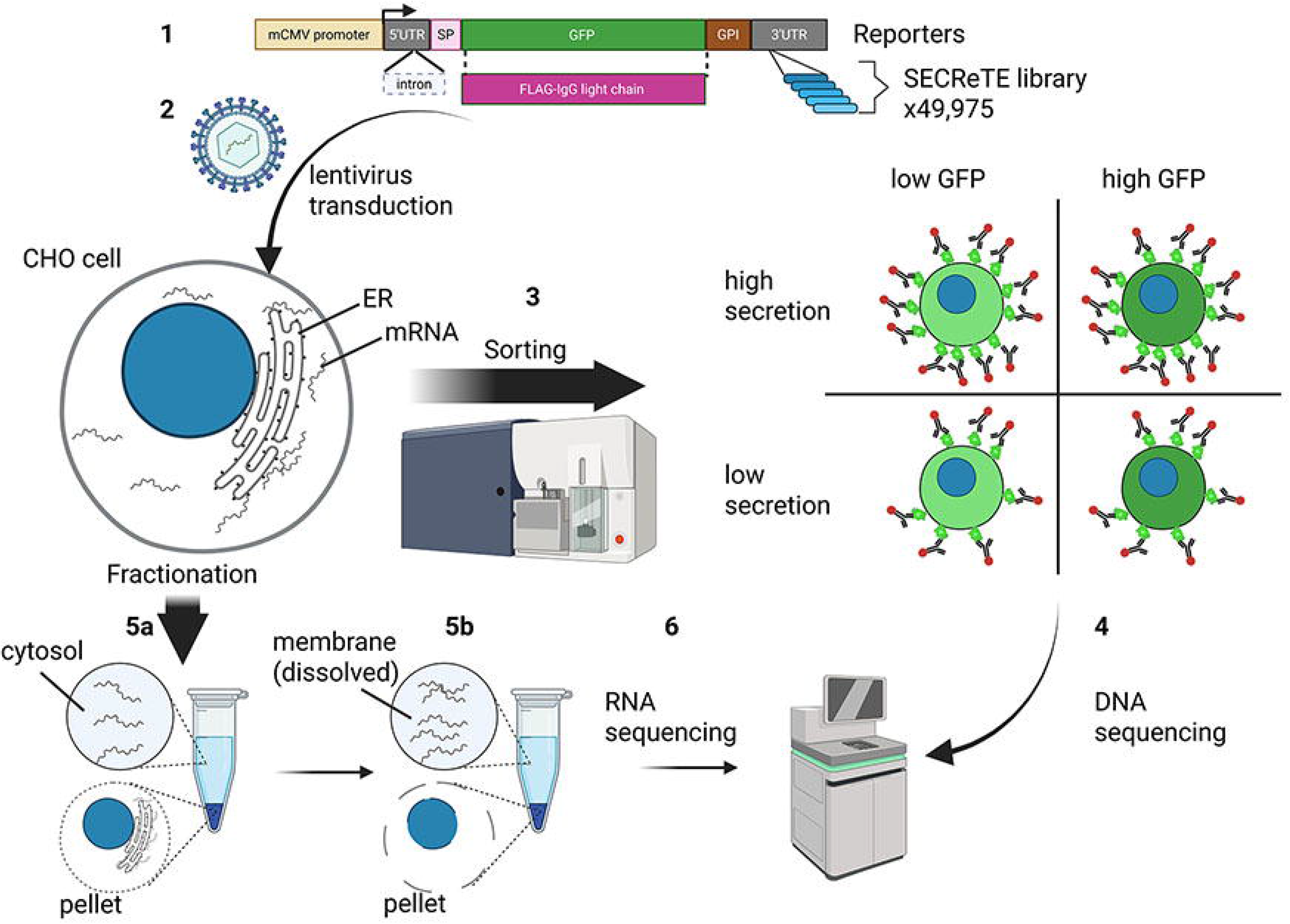
Schematic overview of our high-throughput massively parallel screening of ∼50,000 SECReTE variants. **Step 1**: construction of two reporters expressing GFP or FLAG-LC with a signal-peptide (SP) at the N-terminus and a GPI anchor at the C-terminus. The SECReTE library is inserted into the 3’UTR. **Step 2**: The library is packaged in lentivirus and is used to transduce CHO cells. **Step 3**: GFP library transduced CHO cells are sorted based on GFP expression level and detection of surface GFP (secretion). The LC library was sorted based only on secretion (not shown). **Step 4**: DNA is extracted from the selected populations, and the library region is sequenced. **Step 5a-b**: In parallel, library-transduced CHO cells are fractionated to cytosol and membrane fractions and RNA is isolated from each fraction. **Step 6**: The library region in the mRNAs is sequenced.

## Methods

### Library design

To identify the entire human secretome compendium, we integrated data from the Human Protein Atlas version 18.1 (26), www.proteinatlas.org), Compartments V4.11 and Uniprot databases (27). From the “Human Protein Atlas”, we downloaded a list of pre-computed “secreted” and “plasma membrane” genes. We further analyzed the “cellular location” information that is provided by Uniprot and selected genes that have one of the following keywords in their cellular location: secreted, secretory vesicle, extracellular (for secreted proteins); and cell membrane, GPI anchor, plasma membrane, and cell-surface (for plasma membrane proteins). Cellular compartment information was downloaded via the GeneCards database (28, 29), from which we selected genes having a cellular protein location at the ER, plasma membrane or extracellular. We defined a gene product as “secreted” if it was predicted to be secreted by at least two of the databases. We also added to the list 119 additional genes for which a signal peptide was identified (by Uniprot) and that were predicted to encode secreted/plasma membrane proteins by only one dataset. In total our secretome compendium includes 3127 plasma membrane and 2651 secreted protein-encoding genes (**File S1**). Next, we searched for SECReTE motifs in the secretome using refseq canonical mRNA sequences (by Uniprot). We used the tool that was described in Cohen-Zontag *et al.* (2019) (19) and extracted all SECReTE sequences from these genes. The sequences were then extended from the 5’ and 3’ ends of each SECReTE motif, to yield 26,557 sequences of a total length of 120 bp each (**Files S1** and **S2** and **Table 1**). The distribution of the motifs based on physical and functional position is presented in **Fig S1**.

**Table 1:** List of sub-libraries.

| Sub-library | N | [%] |
| --- | --- | --- |
| **Naturally occurring motifs in human genome** | 26557 | 55.3 |
| **Naturally occurring motifs with fixed flanking sequence** | 4554 | 9.5 |
| **Tiling of chosen secreted proteins** | 3915 | 8.2 |
| **Perturbations on selected SECRéTE motifs** | 5725 | 11.9 |
| **Tandem motifs** | 824 | 1.7 |
| **Randomly chosen motifs** | 6400 | 13.3 |
| **Total** | 47975 |  |

N = number of variants. % = percent from total library

To test whether the effect of the SECReTE motif is dependent on its flanking sequence we randomly chose 4,554 motifs out of the 26,557 motifs found in human genes encoding secretome proteins and generated 120nt sequences where these motifs are placed between constant 3’ and 5’ flanking sequences (**File S2** and **Table 1**).

To explore the existence of secretion-enhancing motifs in secreted proteins in an unbiased fashion, including proteins that lack obvious secretion elements such as signal peptide (SP) or transmembrane domains (TMD), and their mRNAs lack SECReTE motifs, we used a “tiling approach”. For this purpose, we chose genes of 10 secreted proteins (5 of which without SECReTE, SP or TMD) from the human genome, 4 secreted proteins from the Chinese hamster genome, both IgG light and heavy chains, and 4 non-secretome genes as negative controls (see **Table 2** for details). For each of these 20 genes, we systematically scanned their sequence and synthesized for the library non-overlapping windows of 120nts starting at every tenth nucleotide. This subset included 3,915 variants (**File S2** and **Table 1**).

**Table 2:**
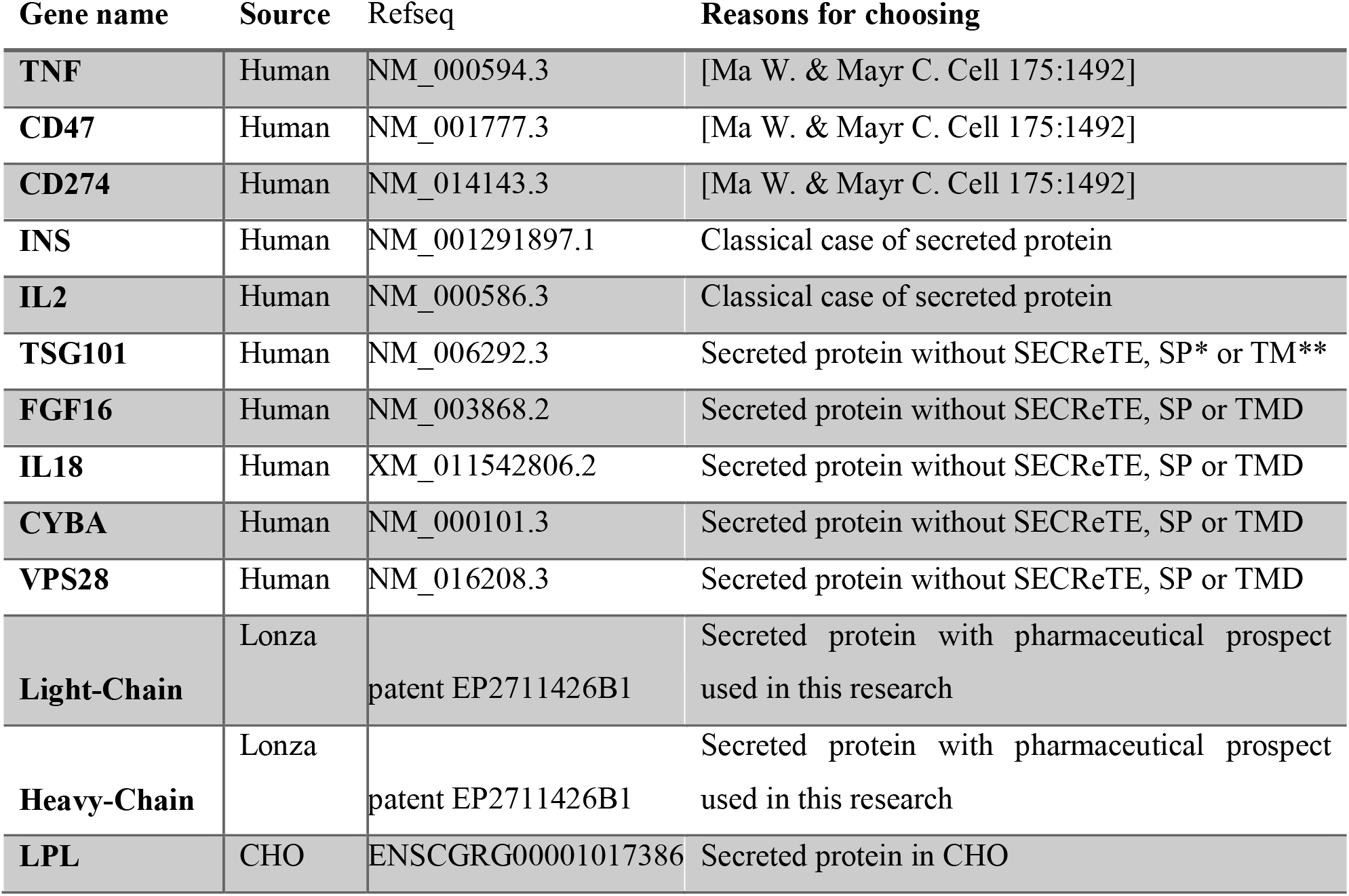
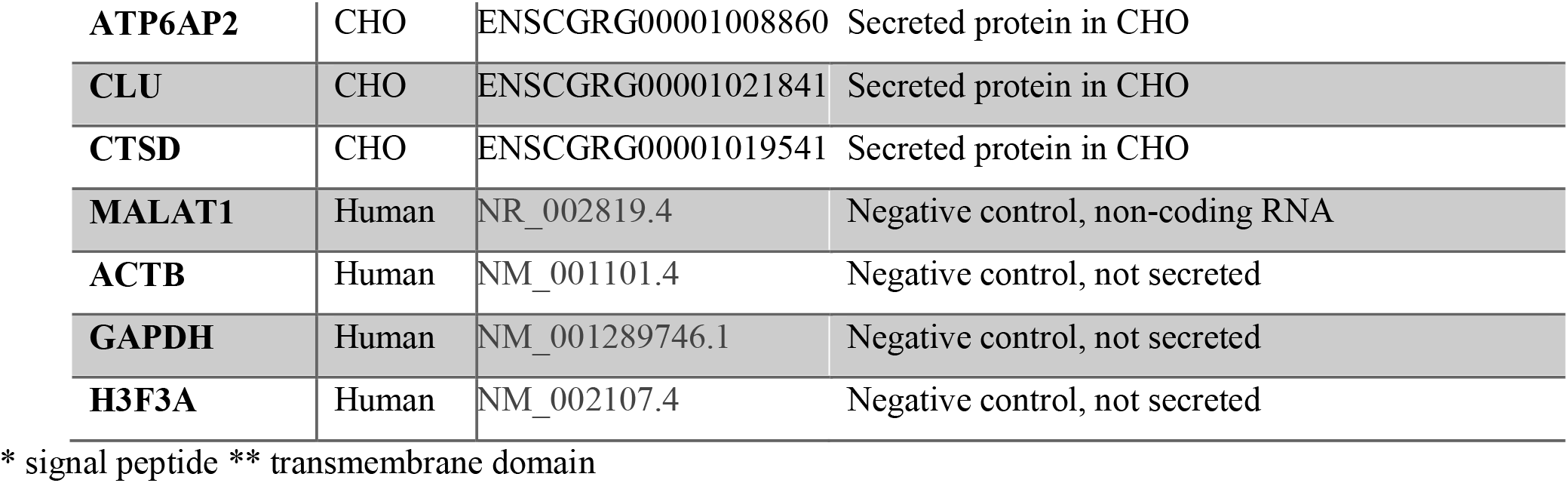
Selected genes for tiling.

| **Gene name** | **Source** | **Refseq** | **Reasons for choosing** |
| --- | --- | --- | --- |
| **TNF** | Human | NM_000594.3 | [Ma W. & Mayr C. Cell 175:1492] |
| **CD47** | Human | NM_001777.3 | [Ma W. & Mayr C. Cell 175:1492] |
| **CD274** | Human | NM_014143.3 | [Ma W. & Mayr C. Cell 175:1492] |
| **INS** | Human | NM_001291897.1 | Classical case of secreted protein |
| **IL2** | Human | NM_000586.3 | Classical case of secreted protein |
| **TSG101** | Human | NM_006292.3 | Secreted protein without SECRETE, SP* or TM** |
| **FGF16** | Human | NM_003868.2 | Secreted protein without SECRETE, SP or TMD |
| **IL18** | Human | XM_011542806.2 | Secreted protein without SECRETE, SP or TMD |
| **CYBA** | Human | NM_000101.3 | Secreted protein without SECRETE, SP or TMD |
| **VPS28** | Human | NM_016208.3 | Secreted protein without SECRETE, SP or TMD |
| **Light-Chain** | Lonza | patent EP2711426B1 | Secreted protein with pharmaceutical prospect used in this research |
| **Heavy-Chain** | Lonza | patent EP2711426B1 | Secreted protein with pharmaceutical prospect used in this research |
| **LPL** | CHO | ENSCGRG00001017386 | Secreted protein in CHO |
| **ATP6AP2** | CHO | ENSCGRG00001008860 | Secreted protein in CHO |
| **CLU** | CHO | ENSCGRG00001021841 | Secreted protein in CHO |
| **CTSD** | CHO | ENSCGRG00001019541 | Secreted protein in CHO |
| **MALAT1** | Human | NR_002819.4 | Negative control, non-coding RNA |
| **ACTB** | Human | NM_001101.4 | Negative control, not secreted |
| **GAPDH** | Human | NM_001289746.1 | Negative control, not secreted |
| **H3F3A** | Human | NM_002107.4 | Negative control, not secreted |

\* signal peptide \*\* transmembrane domain

To test the importance of specific nucleotides in naturally occurring SECReTE motifs we created two mutant sub-libraries. We manually selected 13 SECReTE motifs from the human genome (see **Table 3** for details). Since we used Chinese hamster ovary (CHO) cells for our screen, we selected additional 4 motifs from the Chinese hamster (*Cricetulus griseus*) genome. We also selected the 6 motifs found in the light and heavy chains of the mouse-human chimeric monoclonal antibody (mAb) cB72.3 (30) (provided by Lonza Biologics; sequences are available at Lonza Biologics patent EP2711426B1), since we used this light chain as one of our reporters. For each of these motifs, we introduced all possible single-nucleotide substitutions within the motif sequence. As for the wild-type motifs, these mutated motifs were taken with their natural 3’ and 5’ flanking sequences to complete the 120nt oligo. Since identification of single-nucleotide substitution is sensitive to sequencing errors for this set of library variants, we added a unique barcode as a sequence of 10 randomly chosen nucleotides that replaced the first 10 nucleotides of the oligo (*i.e.* replacing 10 nucleotides from the 5’ flanking sequence). To identify regions in the motif that are important for secretion we used a sliding window approach for mutating the motifs. Meaning that in each such variant a region of 5 nucleotides within the motif was replaced by a random sequence and using a sliding window approach we covered all possible 5 nucleotide regions in all the 17 perturbed motifs. In total, this subset included 4,986 single nucleotide substitution variants and 739 sliding window variants (**File S2** and **Tables 1 and 3**).

**Table 3:** Genes and motifs selected for the motif perturbation sub-library:

| Gene name | Source | # of motifs | SECRete lengths | Reasons for choosing |
| --- | --- | --- | --- | --- |
| **TNF** | Human | 3 | [39,39,33] | [Ma W. & Mayr C. Cell 175:1492] |
| **CD274** | Human | 2 | [36,36] | [Ma W. & Mayr C. Cell 175:1492] |
| **INS** | Human | 1 | 30 | Classical case of secreted protein |
| **IL2** | Human | 1 | 36 | Classical case of secreted protein |
| **SCIN** | Human | 1 | 30 | Secreted protein without SP or TMD, single SECRETE |
| **TIGD7** | Human | 1 | 33 | Secreted protein without SP or TMD, single SECRETE |
| **PDCD10** | Human | 1 | 30 | Secreted protein without SP or TMD, single SECRETE |
| **AVEN** | Human | 1 | 39 | Secreted protein without SP or TMD, single SECRETE |
| **APCS** | Human | 1 | 60 | Single long SECRETE |
| **BMP10** | Human | 1 | 60 | Single long SECRETE |
| **CLU** | CHO | 1 | 30 | Secreted protein in CHO, low motif count |
| **Lgals3bp** | CHO | 1 | 30 | Secreted protein in CHO, low motif count |
| **PSAP** | CHO | 2 | [30,30] | Secreted protein in CHO, low motif count |
| **antibody HC** | Lonza | 4 | [30,36,33,30] | Motifs from the antibody used in this research |
| **antibody LC** | Lonza | 2 | [42,39] | Motifs from the antibody used in this research |

To test whether the effect of dual or triple SECReTE motifs in the same 3’UTR, we chose 16 SECReTE motifs from the human genome and 4 additional motifs from the CHO genome, and created dual or triple combinations between these 20 motifs. For each combination, we created one variant each where the two motifs were separated by a 20nt sequence and one variant each where the two motifs were joined to form a single long motif. The spacing sequence and flanking sequences for this subset were the same randomly chosen sequences for all variants in this subset. Additionally, for 12 out of the 20 chosen motifs that were short enough to fit 3 times in a 120nt oligo sequence we also created variants with 3 repeats of the same motif, once when each of the repeats is separated by a spacing sequence, and once when all three repeats are joined together to create one long motif (**File S2** and **Table 1**).

To explore synthetic SECReTE motifs, we synthesized a set of 2,000 sequences with 39 repeats of an *NNY* triplet. To test whether the wildcard nucleotides (*N*) in the SECReTE pattern can be any nucleotide or if there is a preference for purines (*R*) in these positions, we also generated 2,000 random sequences with an *RRY* pattern. To test the functionality of pyrimidine-rich motifs, we generated 2,000 random pyrimidine-rich sequences with varying pyrimidine frequencies (*e.g.* 70%, 80%, 90% or 100% pyrimidines; 500 sequences each), and as a negative control we generated 400 random purine-rich sequences with varying purine frequencies (*e.g.* 70%, 80%, 90% or 100% pyrimidines; 100 sequences each). For all randomly generated motifs, we avoided low complexity regions by avoiding more than 4 consecutive repeats of the same nucleotide (**File S2** and **Tables 1 and 4**).

**Table 4:** Composition of the random motif sub-libraries.

| Type | Y/R frequency | N |
| --- | --- | --- |
| **polyY** | [0.7,0.8,0.9,1]** | 2000 |
| **polyR** | [0.7,0.8,0.9,1]** | 400 |
| **NNY** | - | 2000 |
| **RRY** | - | 2000 |
| **Total** |  | 6400 |

\*\* - For both the polyY & polyR we created oligos with different Y or R frequency, 25% for each of the listed frequencies.

### Reporter cloning

For cloning, four gene fragments necessary to create the SP-GFP-GPI reporter were ordered from Twist Bioscience and cloned into a pHAGE-UBC-GFP lentiviral vector (Gift from Robert Singer, Albert Einstein College of Medicine, NY) by Gibson cloning. Gene fragments without the signal peptide and GPI were also ordered from Twist and a cytoplasmic GFP reporter vector was created. The CMV promoter, 5’UTR, intron, signal-peptide and 3’UTR sequences were obtained from Lonza Biologics patent #EP2711426B1. The GFP sequence was mutated to destroy an endogenous SECReTE motif in GFP (the codons for V^150^ and I^152^ were changed from *GTC* and *ATC* to *GTG* and *ATA*, respectively). The GPI anchor sequence was taken from human CD59. *Bam*HI and *Mlu*I restriction sites, separated by 8nts, were introduced into the 3’UTR sequence to facilitate library cloning. To alleviate a problem of plasmid toxicity in *E. coli*, the expression cassette was flipped using restriction enzymes *Eco*RI and *Asc*I, so that the entire cassette is on the lentivirus negative strand. The GFP reporter is depicted in **Fig 1, step 1.**

To construct the LC reporter, the LC sequence of mAb cB72.3 (30) was PCR amplified and cloned by Gibson cloning in place of GFP. In this case, the SECReTE motif in the LC CDS was maintained. FLAG tag was introduced after the SP to allow detection by anti-FLAG antibody after SP cleavage. The LC reporter is depicted in **Fig 1, step 1.** To further alleviate bacterial toxicity, both reporter plasmids and libraries were propagated in STBL-3 *E. coli* strain.

### Library synthesis and cloning

Library variants were synthesized as a 120nt ssDNA oligo library by Twist Bioscience. The library was cloned by Twist Bioscience into the lentiviral GFP reporter plasmid between the *Bam*HI and *Mlu*I cloning sites in the 3’UTR. The library plasmids were extracted from the bacterial culture using a midiprep kit (NucleoBond Xtra Midi; Macherey-Nagel). The LC library was cloned in two steps. First, the library oligos were PCR-amplified with primers that included the 5’ *Bam*HI site and 3’ *Mlu*I and *Xba*I sites. Oligos were cloned into the *Bam*HI-*Xba*I sites in pcDNA3.1 vector. Approximately 170,000 colonies were collected and the pcDNA3.1-library plasmids were extracted using the midiprep kit. The library was then sub-cloned into the lentiviral LC reporter plasmid using BamHI-MluI restriction sites. ∼150,000 colonies were collected, and the LC-library plasmids were extracted by the midiprep kit. It should be noted that for maximum yield of the midiprep, 200ml of *E. coli* culture were grown, and plasmid DNA extraction was done with twice the volume of midiprep buffers.

### Packaging into lentivirus

Packaging was performed in HEK293SF-PacLV cells (31) (herein called PacLV cells, a gift from Bernard Massie, McGill University) according to their provided protocol with minor changes. We chose these cells because production is consistent and it is simple to scale-up production. Additionally, PacLV cells contain the three genes required for lentivirus packaging under the control of doxycycline- and cumate-inducible promoters. Thus, there is no need to transfect cells with the three plasmids encoding these genes together with the reporter vector. Briefly, PacLV cells were routinely grown in suspension in polycarbonate shaker flasks with filtered caps in HyCell TransFx-H media (Hyclone) supplemented with 4mM glutamine, 0.1% Kolliphor P188 and 1mg/ml penicillin/streptomycin. Cells were incubated at 37°C, 5% CO_2_ with agitation at 110rpm. For lentivirus packaging, cells were transferred to SFM4-Transfx-293 medium (Hyclone) supplemented with 4mM glutamine and 1mg/ml penicillin/streptomycin. Cells were grown to a concentration of 1.5×10^6^ cells/ml and transfected with 2.67µg plasmid DNA per 1×10^7^ cells using 2µl of PEI-pro transfection reagent (Polyplus) per 1µg DNA. Lentivirus production was induced by adding 1µg/mL doxycycline and 50µg/mL of cumate at 4-6hrs post-transfection and adding 7mM sodium butyrate at 16-24hrs post-transfection. Media was collected 48 and 72hrs post transfection, filtered using 0.45µm membrane, and concentrated by Lenti-X concentrator (Clontech), which was added into the collected media, centrifuged at top speed and resuspended in CD-CHO medium. Lentiviruses were then used immediately for transduction or frozen at -80°C. 10% fetal bovine serum was added prior to freezing.

### CHO cell transduction

CHOK1-SV cells (Lonza Biologics) were cultured in suspension in CD-CHO medium (Thermofisher) supplemented with 6mM glutamine and 1mg/ml penicillin/streptomycin. Cells were incubated at 37°C, 5% CO_2_ with agitation at 110rpm. Transduction to CHO cells was performed at 1×10^6^/ml in 5ml (for small scale, i.e. reporter alone) or 50ml (for the libraries) with lentivirus concentrated from 10-20ml (for small scale) or 300ml (for the libraries) of PacLV cultures. The CHO media was complemented with 6µg/ml polybrene prior to addition of the virus. After 24 hrs, CD-CHO was added to increase the volume 4-fold.

### CHO cell fractionation

The protocol is based on two publications (32, 33). Briefly, 100µg/ml cycloheximide (CHX) was added to the CHO culture 3min prior to harvesting the cells. Next, 2×10^7^ cells were collected by centrifugation (300x*g* for 4min at 4°C) for the subcellular fractionation samples (*i.e.* cytoplasm; Cyto and membrane fraction; Memb) and 1×10^7^ cells for the total lysate (TL) samples. Cells were resuspended in 5ml cold PBS+CHX and kept on ice for 10 min. Cells were collected by centrifugation (1000xg for 4min at 4°C), washed once in 1ml PBS+CHX and resuspended in digitonin-containing 0.5ml Lysis Buffer A (110 mM potassium acetate, 25mM HEPES pH 7.2, 2.5mM magnesium acetate, 1mM EGTA, 0.03% digitonin, 1mM DTT, 100µg/ml CHX, 1mM PMSF, 1x protease inhibitor cocktail, 40 U/ml RNasin) (for fractionation) or Lysis Buffer B (400 mM potassium acetate, 25 mM HEPES pH 7.2, 15 mM magnesium acetate, 1% NP-40, 0.5% deoxycholic acid, 1mM DTT, 100µg/ml CHX, 1mM PMSF, 1x protease inhibitor cocktail, 40 U/ml RNasin) (for TL). Cells were incubated using an end-over-end rotator for 15min at 4°C and then centrifuged (3000x*g* for 5min at 4°C for the subcellular fractionation sample and 7000x*g* for 10min at 4°C for the TL). The supernatants (Cyto, TL) were collected, divided into protein and RNA extraction tubes, and frozen. The Memb sample (pellet) was washed (x2) with Wash Buffer (110 mM potassium acetate, 25mM HEPES pH 7.2, 2.5mM magnesium acetate, 1mM EGTA, 0.004% digitonin, 1mM DTT, 100µg/ml CHX, 1mM PMSF, 1x protease inhibitor cocktail, 40 U/ml RNasin), resuspended in 0.5ml Lysis Buffer B, and processed like the TL sample. RNA was extracted as follows: 1ml of Tri reagent (Sigma) was added and samples were incubated for 10min at room temperature (RT). Then, 200µl chloroform were added, the samples vortexed and transferred to 5PRIME phase lock tubes, and centrifuged at max speed for 15min at 4°C. The upper phase was transferred to a new tube, the RNA was precipitated with an equal volume of isopropanol, washed with 75% ethanol, and resuspended in 50µl water. cDNA was prepared using Quanta Flex cDNA kit (Quanta 95049-100-2). To validate fractionation based on mRNA distribution, semi-quantitative PCR was performed for PABPC1 and GRP94. The protein samples were quantified using the BCA protein assay kit (Pierce) and analyzed by Western blot with Rabbit anti-CANX (OWL) (an ER marker), mouse anti-Tomm20 (Abcam) (a mitochondrial marker), and Rabbit anti-ERK (a gift from Ari Elson lab; Weizmann Institute) (a cytoplasmic marker) to validate efficient fractionation. Note that throughout the paper we also refer to the membrane fraction as the ER-containing fraction and the associated RNAs as ER-localized RNAs. However, this fraction also contains mitochondria (**Fig S2**) and probably other membrane-bound organelles. Due to the presence of the signal peptide in both reporters, it is less likely that the reporter/library mRNAs localize to other organelles besides ER, but this crude fractionation does not allow us to completely exclude this possibility.

### CHO cell sorting and FACS analysis

To sort the library based upon GFP-GPI expression and secretion, 50 x 10^6^ CHO cells previously transduced with the library were collected (unless otherwise indicated). Cells were washed once in CD-CHO and incubated for 30min on ice in 2ml CD-CHO with Alexa647-rabbit anti-GFP antibody (A-31852, Thermofisher) at a 1:200 dilution. Cells were then washed once with cold CD-CHO, resuspended in cold CD-CHO, and taken for FACS sorting (FACSAria III, BD) based upon GFP expression and Alexa647 signal (secretion). About 25 x 10^6^ cells were sorted to the various sub-populations and the rest (∼25 x 10^6^ cells) were collected for the TL sample.

To sort the library based upon LC secretion, 50 x 10^6^ CHO cells were collected, washed once in CD-CHO, and incubated for 30min on ice in CD-CHO with mouse anti-FLAG M2 antibody (F3165, Sigma) at 1:200 dilution. Cells were then washed once with cold CD-CHO, resuspended in cold CD-CHO with 1:400 Alexa488-goat anti-mouse IgG antibody, and incubated for 20min on ice. Cells were than washed once with cold CD-CHO, resuspended in cold CD-CHO, and taken for FACS sorting based on Alexa488 signal (secretion). About 25 x 10^6^ cells were sorted to the various sub-populations and the rest (∼25 x 10^6^ cells) were collected for the TL sample. In both cases, untransduced CHO cells and reporter-alone transduced CHO cells were used as controls for gating.

### Imaging and image analysis

CHO cells expressing the GFP-GPI reporter or cytoplasmic GFP were stained with Alexa647-rabbit anti-GFP antibody, as indicated above, and imaged using a Zeiss AxioObserver Z1 DuoLink dual camera imaging system equipped with an Illuminator HXP 120-V light source, Plan Apochromat 100× 1.4 NA oil-immersion objective, and a Hamamatsu Flash 4 sCMOS camera. Ten 0.5μm step *z*-stack images were taken for the live imaging using a motorized XYZ scanning stage 130 × 100 Piezo, and ZEN2 software at 0.130µm per pixel. Images were processed and fluorescence intensity quantified by FIJI (34).

### Sequencing of fractionated RNA and sorted population DNA

RNA was extracted and cDNA prepared, as indicated in the cell fractionation section. Genomic DNA was extracted using the Monarch genomic DNA purification kit (NEB, T3010S). RNA and DNA concentrations were measured with the Qubit RNA BR assay (Thermofisher Q10210) and Qubit dsDNA BR assay (Thermofisher Q32853). Library sequences of both DNA and cDNA were amplified by 15 rounds of PCR using specific oligos. For this step we used all available genomic DNA and 2µg of cDNA. The PCR products were cleaned by AMPure XP SPRI beads (A63881, Beckman Coulter). The PCR product concentration was measured by Qubit dsDNA BR assay. A second PCR reaction was performed with 3ng of the cleaned samples and 15 cycles of amplification to add Illumina unique dual indexes and Illumina adapters required for sequencing. The amplified products were cleaned with AMPure XP SPRI bead with a bead-to-sample volume ratio of 1.5, followed by quantification with Qubit dsDNA HS kit (Q32851, ThermoFisher). The nano-molar of the cleaned samples was estimated by an equation: 1000x sample concentration (in a unit of ng per ul) / 0.629x size of the final dsDNA fragments (in a unit of bp). An equal number of moles of the samples were pooled together. The pool with final concentration of 2nM was sequenced with paired-end reads by Illumina NovaSeq 6000 SP200 kit (v1.5). For endogenous CHO mRNA fractionation, the cDNA was processed for MARS-seq (35) at the Nancy and Stephen Grand Israel National Center for Personalized Medicine (G-INCPM) at the Weizmann Institute of Science.

### Bioinformatics analysis of MARS-seq

Prior to quantifying gene expression, we assembled the 3’ UTR gene structure of the Chinese hamster genome. This step was necessary because the publicly available gene annotation datasets primarily include the coding sequence region, whereas our MARS-seq libraries capture the 3’ ends of the transcriptome (35). We used Cufflinks (36) with the parameters: overlap-radius 200, min-isoform-fraction 0.2 and Ensembl V99 genes as a GTF-guide. Cufflinks was applied to each library separately, and the results were merged using cuffmerge. The final gene model (**File S3**) was used in the downstream analysis. Fastq readalignment (via STAR against criGriChoV2), gene quantification, normalization, and differential expression testing (DEseq2) was performed using the UTAP pipeline (37). Cellular compartment of the protein products encoded by the sequenced mRNAs was determined by homology to their human counterparts using the Compartment data extracted from GeneCards database (28, 29). A gene was considered differentially expressed in the cytosol or the membrane fractions if the base-mean expression was > 10, the FDR adjusted p-value (p-adj) was < 0.05, and the log2 fold change in at least one of the pairwise comparisons was at least 1. GO term analysis was performed using Metascape (38).

### Processing and mapping of sequencing reads of the libraries

Sequencing reads were demultiplexed by bcl2fastq (v2.20, illumina) without barcode mismatches allowed. All oligos are flanked with same sequence at each side. These regions were trimmed out from the reads by Cutadapt (package in python/3.6.3) (39). The trimmed paired-end reads were merged into single reads by PEAR (version 0.9.10) (40), followed by mapping to the full list of our library variants by Bowtie2 (version 2.3.5.1) (41). The alignments were converted from sam to bam formats, sorted, and indexed by SAMtools (version 1.15.1) (42). Based on the alignment results, unmapped reads, duplicated reads, and reads with mapping quality below 0 were removed by the Python function pysam (python/3.6.3). The read numbers of each variantwere determined by python function Counter (from collections in python/3.6.3). The read counts for all samples were then organized into a single table by MATLAB (R2021b).

### Bioinformatics analysis of the libraries

#### Calculation of variant scores

The main source of data was the number of reads for each variant across the various repeats of each experiment. We used this data to calculate the ER localization scores and protein secretion scores for each library variant. The ER localization scores were calculated as the ratio of the read counts of the variant in the membrane fraction to its read counts in the cytoplasmic fraction. The protein secretion score is the ratio of the read count of a variant in the high secretion subpopulation (*i.e.* p8 in GFP samples, “high” in LC samples) to its read counts in the low secretion subpopulation (*i.e.* p5 in GFP samples, “no” in LC samples). Each read count value was normalized to the total number of reads sequenced in a given sample in order to correct for potential variation in the number of mapped reads between samples. Hence, each of our scores was a ratio of ratios. A pseudo count of 1 was added to each count to allow for log-transformation also in the case of zero reads.

A two-proportion Z-test was used to test for each variant between a pair of samples, for both ER score and the protein score, to determine whether the score reflected significant enrichment/depletion. The null hypothesis was that the proportion in the numerator of the score was equal to the proportion in the denominator of the score. For example, in the case of the ER score, the null hypothesis was that the proportion of read count of a variant relative to the total reads sequenced in the membrane fraction was equal to the proportion of its read count relative to the total number of reads sequenced in the cytoplasmic fraction. To correct for multiple hypothesis testing, we used the Benjamini-Hochberg correction for FDR with α = 0.05.

### Enrichment analysis of classified variants

Based on whether variants are significantly enriched, significantly depleted, or not significantly enriched nor depleted, we respectively classified the variants into 3 groups, high, low, and non-significant (NS). We then developed a statistical test procedure (exemplified in **Fig S3**) to compare the scores of all library variants variants between pairs of samples. We made three types of comparisons: (1) Between biological repeats of the same experiment (for both the GFP and LC reporters for the ER RNA localization and protein secretion assays); (2) The ER RNA localization scores between GFP and LC, and, separately the protein secretion scores between GFP and LC; and (3) Between the ER RNA localization scores and protein secretion scores, separately for GFP and for LC. Given each such two samples, we assess the agreement between the tendencies of the variants to show a similar score categories (i.e. high, low or non-significant). All such pairwise comparisons are presented in the form of 3×3 matrices. These matrices report on the size of overlap of sets of variants at high, low, and non-significance of each pair of assays. For example, **Fig S3** reports the agreement between biological repeats of the GFP RNA localization experiment. The set of variants that scored high in the two repeats showed an overlap that is 5.4 times larger than expected by chance (expected value gauged as product of frequencies of the high variants relative to all variants in each assay, multiplied by the total number of variants that were assayed in the two assays. See details in **Fig S3** legend). We also report the hypergeometric p-value on the overlap between these two sets was extremely low (Inf in this computation indicates value <10^-300^). High values on the main diagonal of the matrix (*i.e*. on upper right and lower left entries) indicate a positive correlation between the respective pair of samples.

### Data pre-processing

Utilizing ER scores and protein scores, we filtered out variants whose scores were not significant in at least one of the repeats, separately for ER score and for protein scores. Finally, variants that had significant scores in all repeats but that were inconsistent between repeats (that is, enriched in some but depleted in others) were also filtered out. For GFP, we filtered out variants with more than two-fold variation between the two repeats. For all other variants, the final score for each variant was the mean of the scores obtained in the two repeats. For LC, if the three repeats were similar to each other, that is, the fold-variation between the most different pair of repeats was smaller than 2, then the final score was calculated as the mean of all three repeats. If the fold-variation between the most different pair of repeats was higher than 2, but there existed a pair of repeats whose fold-variation was smaller than two, then the final score was calculated as the average between these two repeats. Finally, if the fold-variation between the most similar pair of repeats was higher than 2, then the corresponding variant was filtered out.

### Motif analysis

Motif analysis was performed using MEME, a *de novo* motif discovery algorithm (https://meme-suite.org/meme/)(43).

## Results

### Subcellular fractionation into an ER-enriched membrane fraction and cytosolic fraction

To help better understand whether natural occurrences of the SECReTE motif present within genes affects mRNA localization to the ER membrane, as posited in a previous study (19), we first employed subcellular mRNA fractionation to separate the content into a membrane fraction enriched with the ER membranes and a cytosolic fraction (see *Methods section*). We validated our subcellular fractionation assay by fractionating CHOK1-SV cells by Western blotting and RT-PCR (**Fig S2**). We examined the enrichment or depletion in the membrane fraction and in the cytosolic fraction of three proteins: Calnexin (CANX), which is an ER protein, ERK, which is a cytosolic protein, and TOMM20, a mitochondrial protein that should localize to the membrane fraction as well. As can be seen, the three markers distribute as expected (**Fig S2A**), indicating that the fractionation was successful. Next, we examined the mRNA levels in the different fractions (by RT-PCR) for the GRP94 and PABPC1 genes, which encode ER and cytosolic proteins, respectively. GRP94 was found mostly in the membrane fraction, whereas PABPC1 distributed equally between both fractions (**Fig S2B**), similar to previously published results (33).

Next, we examined the partition ratio between the membrane and cytosolic fractions for each endogenous CHO cell mRNA tested. To determine their expression profile, we reverse transcribed the mRNA derived from the different fractions and subjected it to MARS-seq (35) (**File S4**). Reassuringly, we found that mRNAs encoding secreted proteins are enriched in the membrane fraction (Membrane; *i.e*. 23.7% of membrane fraction mRNAs encode plasma membrane proteins, 19.4% ER proteins, 11.6% extracellular proteins and 10.5% Golgi proteins) whereas the cytosolic (Cytosol) fraction is enriched with mRNAs encoding cytosolic proteins (30.5%) and nuclear proteins (35.1%) (**Fig 2A**, **File S4**). We also examined the presence of the SECReTE motif among the membrane-enriched mRNAs and found that the motif is enriched among the mRNAs localized in the membrane fraction (mean value of SECReTE score for Membrane 4.01±3.54 *vs*. 1.86±2.61 in the Cytosol; p < 2.2 x10^-16^), as expected (**Fig 2B**, **File S4**). GO term analysis found membrane enrichment of mRNAs encoding secreted proteins (e.g. glycoprotein metabolic process, transporters and extracellular matrix) and depletion of non-secreted proteins (e.g. RNA metabolism, Cell cycle, protein modifications and translation) (**Fig 2C-D**). Thus, our fractionation procedure separates out the membrane-enriched fraction of the cell along with mRNAs enriched with the SECReTE motif.

**Fig 2.**
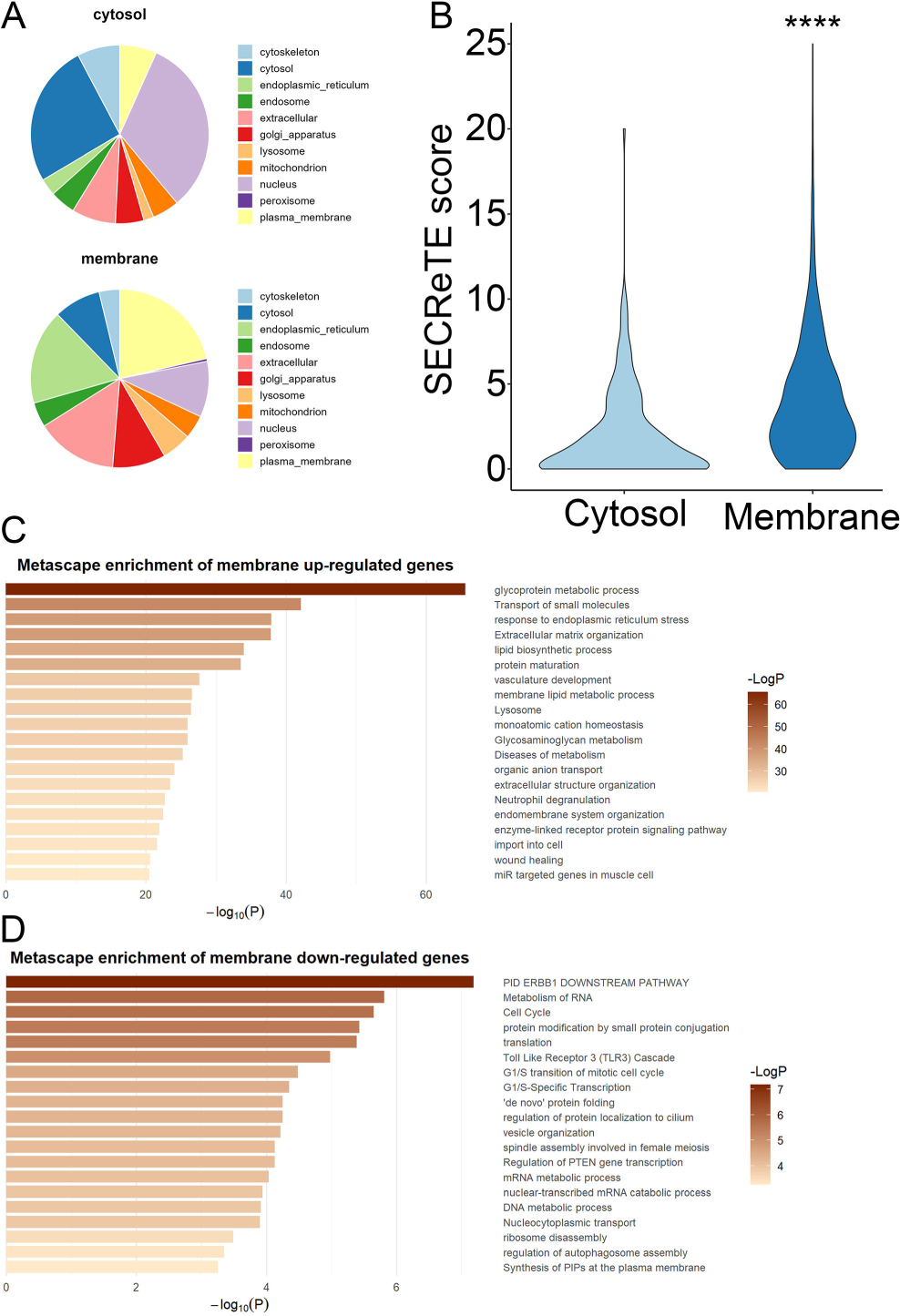
Classification and SECReTE scores of mRNAs in cytoplasm vs. membrane fractions. **A)** Pie charts showing the distribution of compartment classifications of mRNAs in the membrane fraction (top) and cytosolic fraction (bottom) (see also **File S4**). **B)** Violin plot of the SECReTE scores (based upon the number of *NNY* triplet nucleotide repeats) of mRNAs present in the cytosolic or membrane fractions (see also **File S4**). **\*\*\*\*** - p < 2.2×10^-16^. **C-D)** GO term analysis of membrane up-regulated (**C**) and down-regulated (**D**) genes.

### Validation of the GFP-GPI and LC-GPI reporters

Two reporters were designed for library screening (**Fig 1, step 1**). Both reporters enable the protein products to enter the secretory pathway using a signal peptide and be secreted, but also to remain on the cell surface due to the presence of a GPI anchor. This allows for systematic and quantifiable detection of secretion associated with each variant in the library. To maintain consistency between reporter gene expression, we used the same promoter, 5’ and 3’ UTRs, and signal peptide (SP), the latter which notably harbors a SECReTE motif within its encoded hydrophobic residues (*15x NYN*).

The first reporter was based on the green fluorescent protein, GFP. This reporter allows for the quantification of both the protein expression level and level of secretion by FACS analysis (**Fig 1, step 3**). For the second reporter, we chose the IgG Light Chain (LC) gene. Note that the LC gene harbors an additional SECReTE motif (*13x NYN*) in its CDS. This reporter allows for the quantification of only the level of secretion (high or low) by FACS analysis. We also created a control cytoplasmic GFP reporter which lacks either the SP or GPI anchor (cytoGFP).

We first created CHO cells expressing either the GFP-GPI reporter or the cytoGFP control and checked their expression and localization by microscopy. The images show that extracellular GFP-GPI is recognized by the anti-GFP antibody added to the live cells, but not the cell-internal GFP or cytoGFP, as expected (**Fig 3A-B**). We note that the expression level of the cytoGFP is ∼30-fold higher compared to the GFP-GPI reporter (**Fig 3C**). Similar results were obtained when comparing the GFP expression level by FACS analysis (**Fig 3D**). Cell fractionation analysis shows that the GFP-GPI mRNA is found mainly in the membrane fraction, similar to GRP94 mRNA, whereas cytoGFP mRNA is equally distributed between fractions, similar to PABPC1 mRNA (**Fig 3E**). This is the expected result since the signal peptide should tether the mRNA to the ER during translation. We also noticed that the percentage of GFP-positive cells declines within days after transduction with the GFP-GPI (results not shown). The reason for this is not known, although it may relate to toxicity of the membrane-anchored reporter, perhaps by interfering with normal cellular functions. Unavoidably, this situation was found to affect library screening and analysis (see below). For the LC-GPI reporter, we tested our ability to detect the protein on the cell surface by FACS analysis. Indeed, cell surface staining was detected for CHO cells transduced with the LC-GPI reporter and only low background staining of the FLAG antibody was detected for untransduced CHO cells (**Fig S4**).

**Fig 3.**
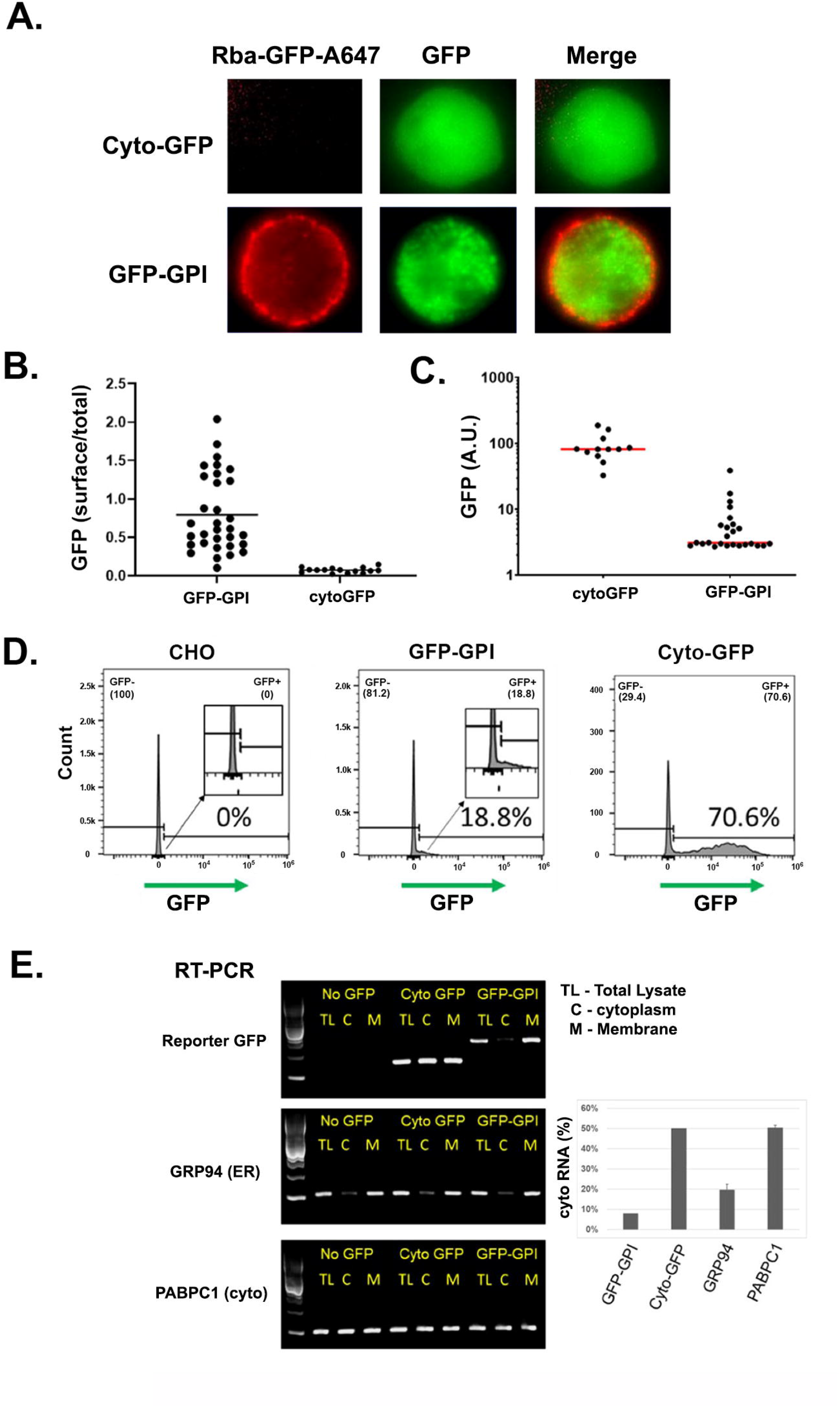
Expression and localization of GFP-GPI protein and mRNA. **A)** Microscope images in the GFP channel and far-red (Alexa647 (A647) Rabbit anti-GFP antibody) channel show the localization of the two GFP reporter constructs. Note: exposure time in the GFP channel for the GFP-GPI was longer than for cytoGFP, due to differences in expression levels. **B)** Quantification of the ratio of Alexa647 anti-GFP signal intensity over total GFP signal intensity. Each dot in the plot is an individual cell. **C)** Quantification of the GFP expression levels when the exposure time for imaging GFP-GPI and cytoGFP is equal. Each dot in the plot is an individual cell. **D)** FACS analysis of CHO cells alone or CHO cells expressing cytoGFP or GFP-GPI. The indicated percentage is % GFP-positive cells. X-axis is GFP fluorescence intensity. **E)** Cell fractionation followed by RT-PCR of the indicated mRNAs shows the distribution of the GFP-GPI and cyto-GFP mRNAs. Band intensity was quantified by FIJI and % cytoplasmic mRNA was calculated as the ‘C’ fraction divided by ‘C’+’M’ fractions.

### Design, cloning and validation of the SECReTE variant libraries

We designed a library composed of 47,975 different synthetic variants, including naturally occurring human SECReTE motifs, tiled secreted proteins from human and CHO cells, as well as tandem and randomized variants which were divided into several sub-libraries, as detailed in **Table 1**. The composition of the randomly chosen motifs sub-library is depicted in **Table 2** and those of the selected secreted genes for tiling and motif perturbations are listed in **Table 3** and **Table 4**, respectively.

The library was synthesized and cloned into the GFP reporter plasmid by Twist Bioscience. Their quality control sequencing showed 96.3% coverage of the library (*i.e.* 46193 variants). However, the distribution of the library is uneven, with 8552 variants (17.8%) having only 1-9 reads (**File S5**). Moreover, transformation of the plasmid library into STBL-3 cells (followed by plasmid midiprep) reduced the library complexity to ∼28,200 variants, with a similar uneven distribution (**Fig S5A, blue; File S6**). This uneven distribution (also apparent in the original Twist’s sequencing data, not shown) was not due to an effect of homopolymer nucleotides or CG repeats (**Fig S5B-C**), and filtering out homopolymers and CG repeats did not change the distribution (**Fig S5A, orange)**. In parallel, we also sub-cloned the library into the LC reporter and found a similar level of coverage (24,875 variants) and distribution (**Fig S5D**).

### High throughput sequence measurements of GFP and LC reporter genes containing SECReTE variants

We performed two replicates of the GFP library screen and three replicates of the LC library screen. To achieve the maximum percentage of reporter-positive cells (due to the decline in GFP and LC expression over time), the libraries were sorted or fractionated between 2 to 3 days post-transduction. We collected four populations from each of the two GFP-GPI libraries (**Fig 1, step 4** and **Fig S6**) and three populations from each of the three LC-GPI libraries (**Fig S6**), as well as a sample of a whole, unsorted library from each culture. We collected genomic DNA and amplified the 3’UTR region of the library by PCR using oligonucleotides specific to the SECReTE variant-containing region. DNA sequencing allowed us to quantify the variants present in each sorted sub-population. In addition, we fractionated cells from two GFP-GPI and three LC-GPI cultures and collected mRNA from the cytosolic and membrane fractions, as well as from unfractionated cells (total), followed by RT-PCR amplification of the 3’UTR. RNA sequencing allowed us to quantify the variants of the reporter mRNAs enriched in each of the sub-cellular fractions. Sequencing of the resulting genomic DNA and the cDNA samples revealed a high level of coverage of the libraries (*i.e.* 25,000-30,000 variants for the GFP samples, and 23,000-28,000 variants for the LC libraries with at least 1 read, and >20,000 variants with ≥10 reads for most of the samples) (**Fig S7**). Total read count varied between samples from 5.9 to 41 million reads, but most samples had between 10 to 20 million reads. Read counts of each variant in all samples and total number of reads can be found in **File S7**.

We calculated the scores of every variant in each sample (see *Methods* section). The scores of the 32,330 variants that passed the filtering steps in at least one type of experiment (see data preprocessing in *Methods* section) can be found in **File S8.** We then calculated the Z-Scores of each of these variants and were left with 24,187 variants with a final score in at least one assay (**File S9**). The final number of variants from each sub-library and each type of experiment or calculated score that were used for further bioinformatics analysis is shown in **Table 5**.

**Table 5:**
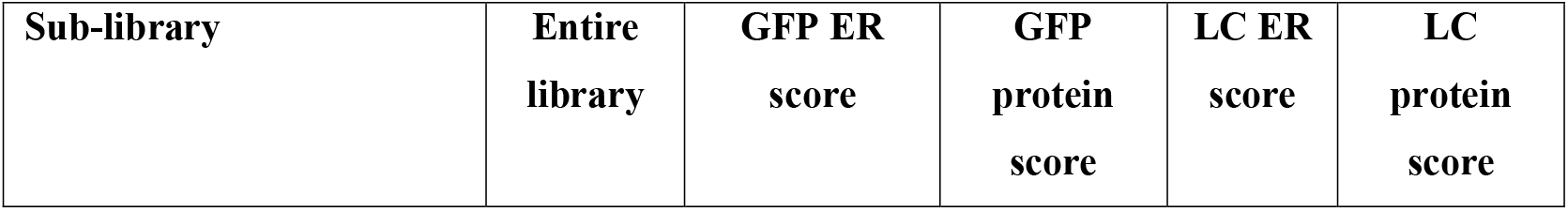
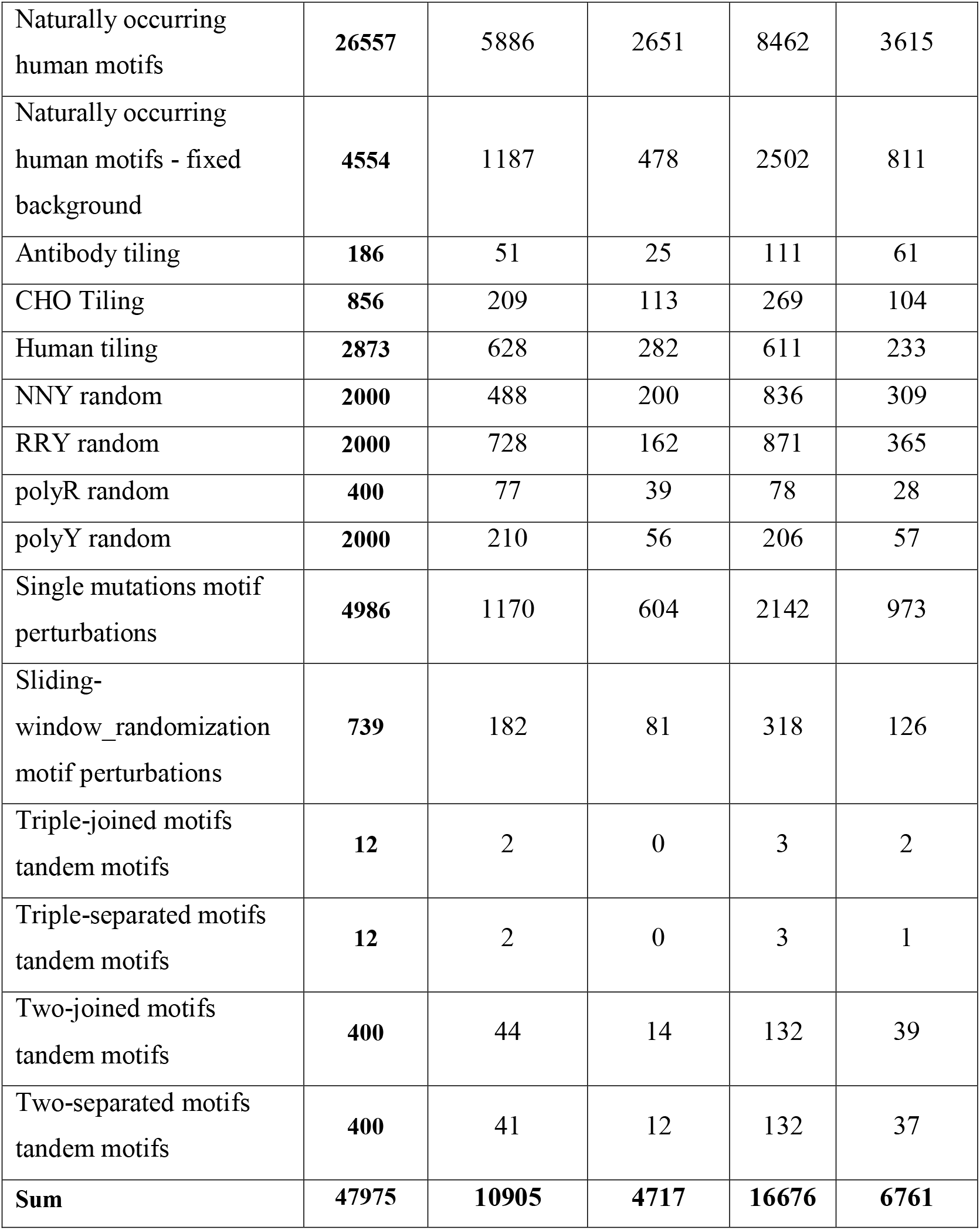
The number of variants of each sub-library and for each score.

### Bioinformatics analysis of the GFP and LC reporter library screens

We plotted the distribution of the ER log_2_(score) for the LC library **(Fig 4A**). The histogram is bimodal since only variants in the high or low ER enrichment categories are shown, as the rest have no significant tendency to be enriched in either the membrane fraction or the cytosol. Most of the variants were enriched in the cytosolic fraction (4852 designs) compared to the membrane fraction (2530 designs). Replicates #1 and #2 showed reproducible results (**Fig 4B**). We then looked at the correlations between all three replicates by calculating the observed-to-expected ratios corresponding to three categories (See **Fig S3** for explanation): 1) enriched in the ER fraction; 2) depleted from the ER fraction; or 3) non-significant (ns). All replicates were highly correlated with each other, as can be seen by the high ratio of observed-to-expected (green numbers) and the -log_10_ p value (red numbers) in the diagonal from the top left to bottom right squares (**Fig 4C**).

**Fig 4.**
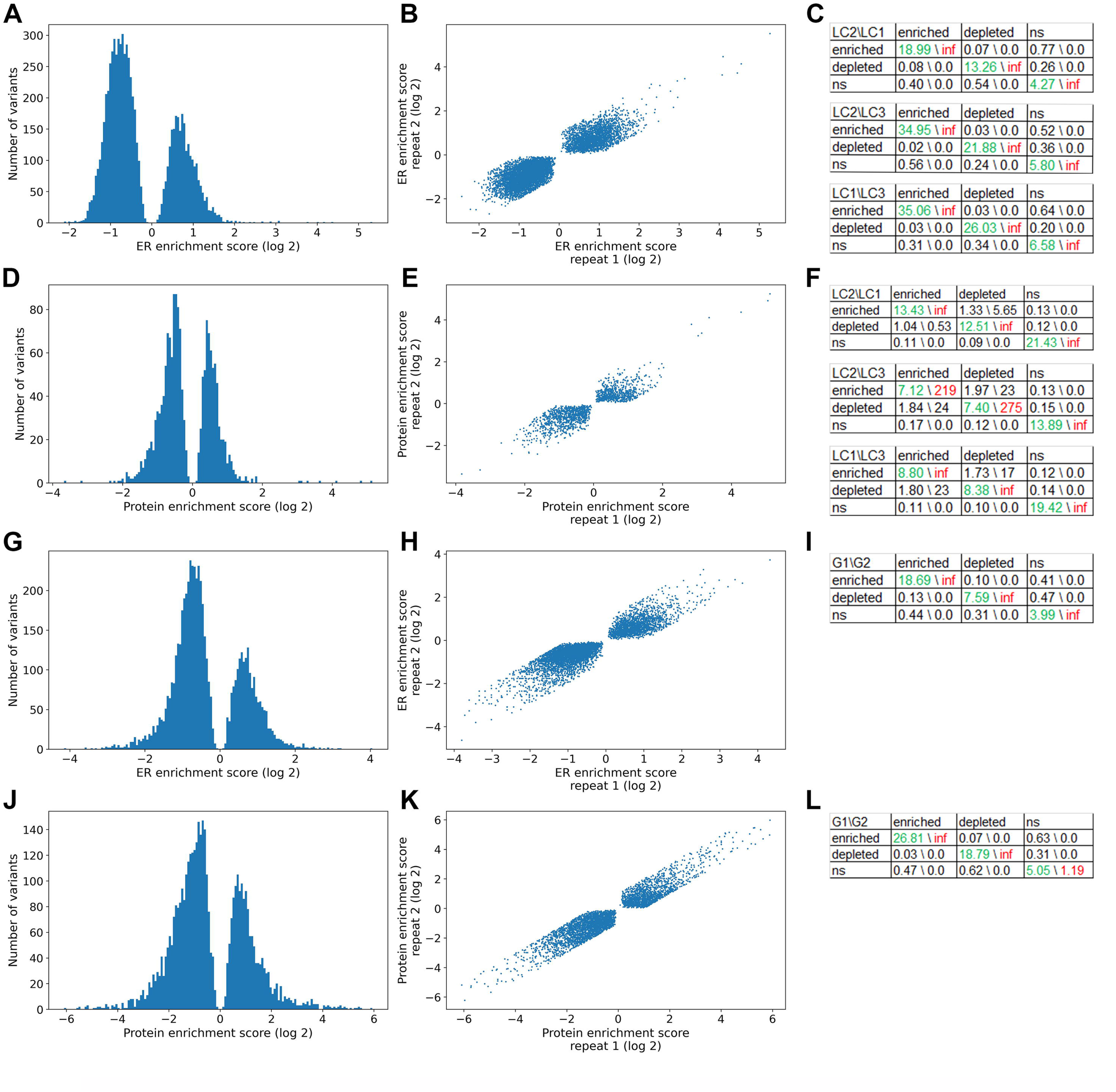
Distribution of scores and agreement between biological repeats. Graphs showing the distribution of log_2_(score) of the variants for each type of reporter and score are shown on the left (LC ER score, LC protein score, GFP ER score and GFP protein score; panels **A, D, G** and **J** respectively). In the middle are graphs showing the agreement between replicates #1 and #2 of the LC reporter (panels **B** and **E**) and replicates #1 and #2 of the GFP reporter (panels **H** and **K**). Each dot is a single variant. On the right are enrichment analysis 3×3 tables of the repeats for ER score (panels **C** and **I**) and protein scores (panels **F** and **L**). Each table compares between two replicates. LC1, LC2, LC3 - LC reporter replicates. G1, G2 - GFP reporter replicates. Variants are classified into high (enriched), low (depleted), or non-significant (ns) changed levels in membrane-localized RNA or secreted protein levels, based on the relative presentations of variants (see Enrichment Analysis in Methods for details and **Fig S3** as an example). Numbers in green are the ratio of the observed to expected number of sequence variants overlapped between 2 categories for indicated 2 samples. Numbers in red, -log_10_ of p-value of statistical significance for the overlaps from Hypergeometric test. In this test “inf” stands for p-value<10^-300^.

We next examined protein secretion from the different library variants. The histogram of secreted fraction enrichment is bimodal too (**Fig 4D**) and the reproducibility across repeats is high in this assay as well (**Fig 4E-F**). For the LC protein score we again see more variants that are depleted (*e.g.,* 655 low secretion variants) than enriched (*e.g.,* 470 high secretion variants) in the secreted fraction.

We then plotted the enrichment values for the GFP library (**Fig 4G-L**). Again, we found that for the ER score, most variants were enriched in the cytosolic fraction (3862 designs) compared to the membrane fraction (1818 designs), and for the protein score most variants are depleted (2593 low secretion variants) than enriched (1523 high secretion variants). As for the LC library, we see a strong correlation between the two repeats for both scores.

### Correlation between RNA association with ER membranes and protein secretion

We next asked if the SECReTE variants associated with high levels of ER membrane localization are also those that are associated with a high degree of protein secretion. **Fig 5A** shows that this is indeed the case for the GFP reporter, *i.e*. there is a highly significant overlap (depicted by *p* value) between the set of variants that scored high for both ER RNA localization and protein secretion for both types of analysis. These results support the basic notion that mRNA association with the ER indeed enhances protein secretion. However, there was little-to-no correlation between the ER and Protein scores for the LC library (**Fig 5B**).

**Fig 5.**
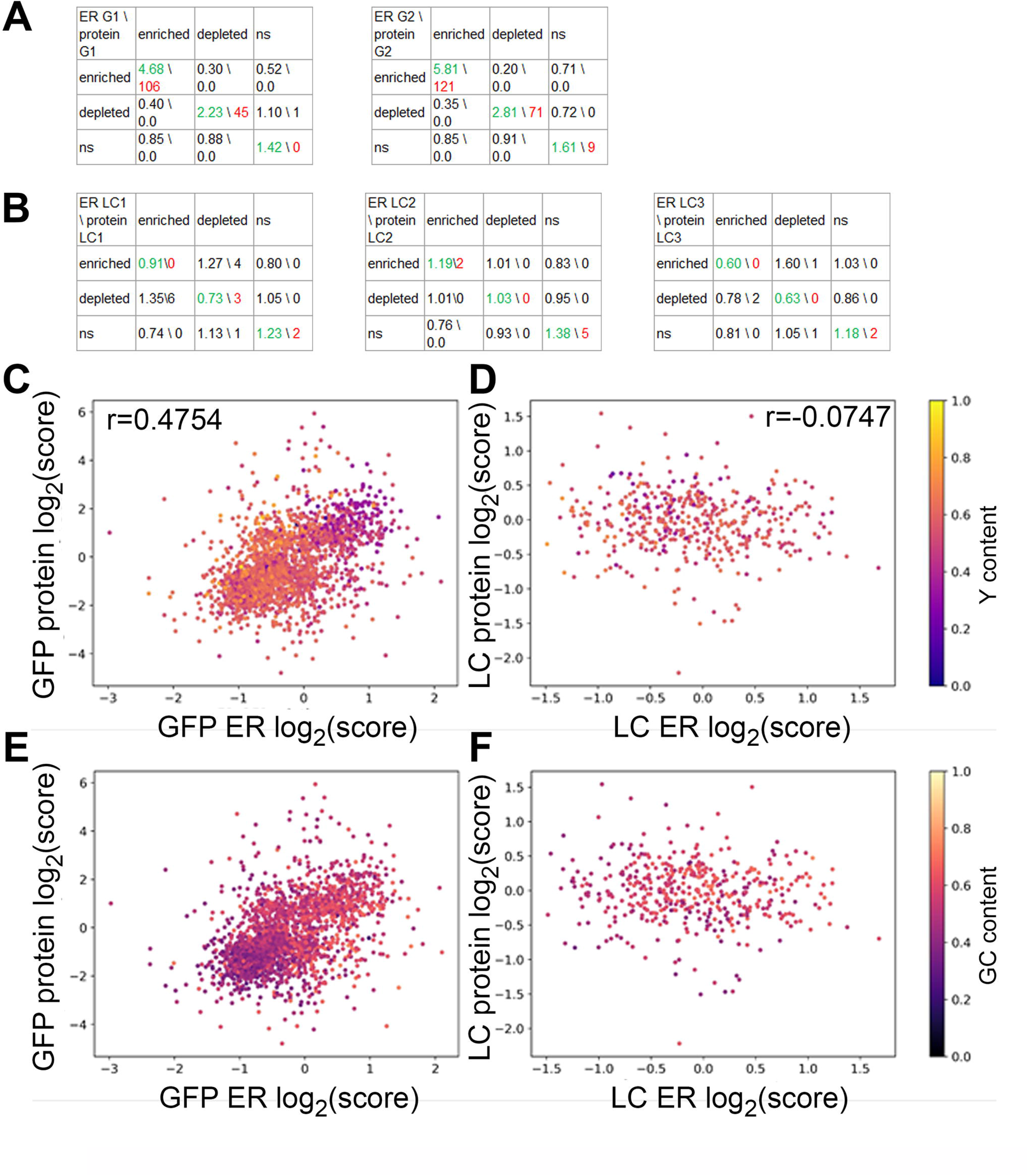
Positive correlation between ER and protein scores. Panels **A** and **B** show enrichment analysis 3×3 tables of ER score *vs* protein score of the various replicates. LC1, LC2, LC3 - LC reporter replicates. G1, G2 - GFP reporter replicates. Variants are classified into high (enriched), low (depleted), or non-significant (ns) changed levels in membrane-localized RNA or secreted protein levels, based on the relative presentations of variants (see Enrichment Analysis in Methods for details and **Fig S3** as an example). Numbers in green are the ratio of the observed to expected number of sequence variants overlapped between 2 categories for indicated 2 samples. Numbers in red, -log_10_ of *p* value of statistical significance for the overlaps from Hypergeometric test. In this test “inf” stands for p-value<10^-300^. (**C-F**) graphs showing ER log_2_(score) (x-axis) *vs* protein log_2_(score) (y-axis) of the GFP library **(C, E)** and LC library **(D, F)**. Each dot is a single variant from the library. Pearson correlation (r) is noted at the top of the graphs. The color of the dots indicates the Y content **(C-D)** or GC content **(E-F)** as fraction of nucleotides. See color bar on the right.

We next asked if RNA association with the ER membrane fraction correlates with protein secretion across variants of the library on a continuous scale, rather than the 3 categories (*i.e.,* enriched, depleted or non-significant (ns)).

In the GFP reporter library we found a significant positive correlation between mRNA association with the ER fraction and protein secretion (**Fig 5C-D**; Pearson correlation r=0.4754; 95% CI (0.4426 to 0.507); p<0.0001), meaning library variants enriched with the ER at the RNA level tend to encode proteins that are secreted better. Variants with high ER and protein scores tended to have low Y content (**Fig 5C**) and somewhat enriched GC content (**Fig 5D**). Interestingly, of the top variants with log_2_(score) >0.5 in both ER and protein scores (219 variants), 37.4% contained natural SECReTE motifs and 35.6% contained random RRY motifs.

In the LC, there was no correlation nor anti-correlation between ER localization and protein localization (**Fig 5E-F**; Pearson correlation r=-0.0747; 95% CI (-0.1 to -0.0494); p<0.0001). Variants with very low Y content tended to have a low ER score, but Y content did not seem to affect the protein secretion score. GC content seemed to be spread evenly, irrespective of ER or protein score.

Next, we compared the variants of the two libraries in each of the two scores. We first performed enrichment analysis and found there is a significant overlap between the set of variants of the GFP reporter and LC reporter that scored high (enriched) or low (depleted) groups for both ER RNA localization (**Fig S8A, left**) and protein secretion (**Fig S8A, right**).

We next looked at the correlation on a continuous scale. For the ER score, we found a significant positive correlation between the GFP and the LC libraries (**Fig S8B**; Pearson correlation r=0.3354; 95% CI (0.3105 to 0.3598); p<0.0001), meaning variants that tended to be ER-enriched with the GFP reporter also tended to be ER-enriched with the LC reporter. However, for the protein scores the correlation between variants in the GFP and LC libraries was weaker, yet still positive (**Fig S8C**; Pearson correlation r=0.1723; 95% CI (0.1165 to 0.2269); p<0.0001).

### Comparison between the sub-libraries

Our library consists of a number of sub-libraries, each with different types of motif/syntax and variations thereof. We compared the scores of each sub-library to each other. For the GFP reporter, variants of the RRY sub-library were almost all enriched with the ER and showed high secretion scores (*i.e.,* positive log_2_(score)) (**Fig 6A-B**), having the highest mean and median for the GFP ER scores and GFP protein scores (see **File S9,** descriptive stats sheet). The RRY sub-library was also superior in the LC protein score, but not the LC ER score (**Fig 6C-D, File S9**). Interestingly, the antibody tiling sub-library had a high mean and median for both reporters and in both assays (**Fig 6A-D, File S9**). The single mutation and sliding window sub-libraries, as well as natural SECReTEs with fixed flanking regions, were highly ER enriched for the LC reporter (**Fig 6C**). In the other sub-libraries, most variants tended to be ER-depleted and with low secretion scores, but some variants were found with high enrichment scores. We performed one-way ANOVA analysis on the sub-libraries and found, as expected, that RRY enrichment was statistically significant against most other sub-libraries and in both reporters and assays (**File S10**).

**Fig 6:**
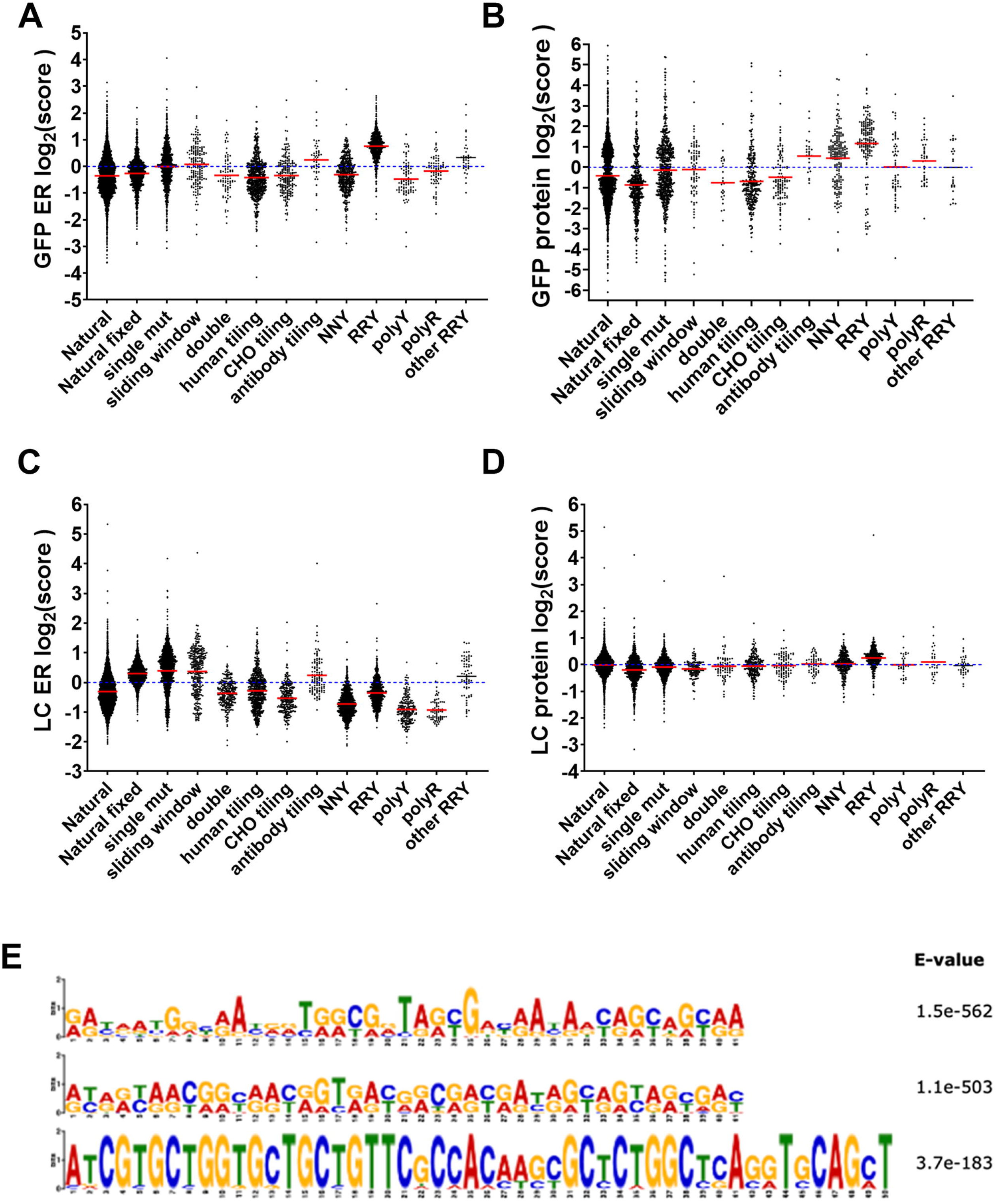
Score distribution of each sub-library and emerging RRY motif. Graphs showing the score distribution of variants from each sub-library. Each dot is a single variant. Red line is the average. The category “other RRY” are natural SECReTEs with an *RRY* motif (see text for more details). See **File S9** for descriptive statistics and **File S10** for one-way ANOVA analysis. **A)** Distribution of the GFP ER score. **B)** Distribution of the GFP protein score. **C)** Distribution of the LC ER score. **D)** Distribution of the LC protein score. **E)** MEME logos for the high-scoring motifs from the GFP ER enriched variants. The two top motifs have an *RRY* repeat structure.

### Motif analysis identifies the *RRY* signature as top scorer for the GFP reporter

We next examined if the top scoring variants carry a distinct signature motif that is more precisely defined than a general *NNY*x10. We performed MEME analysis on the top scoring variants in each assay for ER RNA membrane association and protein secretion and for each of the two reporters (**File S11**). For ER RNA localization with the GFP reporter, the top two scoring motifs have a perfect *NNY*x13 signature, and the third highest motif has a *NNY*x7 motif (**Fig 6E**). Interestingly, the “*N*”s in the *NNY*x13 motifs were invariantly “*R*”, that is purines (*i.e*. *A* or *G*) and never a pyrimidine. Thus, the best scoring motif is actually *RRY* and not *NNY*. For the GFP protein secretion assay, a motif with *NNY*x16 scored highest. The second-highest motif had *NNY*x7 repeats followed by a *R*-rich region. The third motif has a *Y*-rich *NNY*x8 repeat pattern **(Fig S9A)**.

We next searched for motifs from the top scoring variants in the LC reporter assays. The top scoring ER RNA localization motif has a *NNY*x9 repeat motif followed by an *R*-rich region. The second motif was R-rich with no SECReTE-like pattern and the third motif has a *NNY*x10 repeat **(Fig S9B)**. On the other hand, the LC protein secretion score produced three motifs, one of which is a short (20nt-long) sequence composed only of pyrimidines, another has a *NNY*x8 repeat and the third had no SECReTE-like region **(Fig S9C)**. Thus, the most common motif of high scoring variants for both reporters have multiple *NNY* repeats, with a clear preference to *RRY* for GFP ER localization.

Since the *RRY* sub-library had high scores (**Fig 6**), the *RRY* repeat motif discovered by MEME could be the result of a bias due to this sub-library. We therefore searched for *RRY*x7 or longer in the entire library, yet excluding the *RRY* sub-library. We found 114 variants with *RRY*x7 up to *RRY*x40 (**File S9**) that had a score in at least one of the reporters and assays. We plotted these variants as “other *RRY*” against the other sub-libraries (**Fig 6**). Reassuringly, the “other *RRY*” sub-library scored high in both GFP and LC ER scores and somewhat enriched in the GFP protein score. For the GFP and LC ER scores, the “other RRY” group was statistically significantly higher than most other sub-libraries (**File S10**). However, it was not statistically different from the other sub-libraries in both GFP and LC protein scores.

### Analysis of the tiling sub-libraries

The tiling sub-libraries consisted of sequences taken from a selection of secreted genes from human and CHO cells, some of which do not contain SP, TMD or SECReTE motifs. We figured that if these proteins are secreted nonetheless, they might contain other secretion-enhancing motifs that can be discovered in an unbiased way. We synthesized sequence windows based upon tiling the length of these genes, such that each mRNA examined consists of multiple tiles that were scored for ER mRNA localization and protein secretion using the two reporters. The human genes examined in this sub-library are TNF, CD47, CD247, INS, IL2, TSG101, FGF16, IL18, CYBA, and VPS28, and the CHO genes are LPL, ATP6AP2, CTSD and CLU. We also tiled the mRNAs encoding the heavy chain (HC) and light chain (LC) of the mouse-human chimeric monoclonal antibody (mAb) cB72.3. This is the same LC sequence also used as the LC reporter. In addition, we had four negative control genes encoding three non-secreted proteins and one lncRNA (*e.g.* ACTB, GAPDH, H3F3A and MALAT1). We asked if the scores of genes encoding secreted proteins might be different than the controls. The ER scores of most of the secreted proteins tiling variants were low, with a clear tendency for depletion from the ER (negative log_2_(score)) with both reporters **(Fig 7A and C)**. The few secreted proteins with a clear ER enrichment tendency were the l heavy chain, as well as insulin (INS) and VPS28. All three showed statistically significant enrichment in the ER score compared to the others (**File S10**), and HC and VPS28 were also statistically enriched in the LC ER score. Of the negative controls, MALAT1 showed a statistically significant strong depletion tendency compared to HC, INS, GAPDH, VPS28 and ACTB (**File S10**). ACTB and GAPDH seemed to be evenly distributed, with a slight tendency for enrichment.

**Fig 7.**
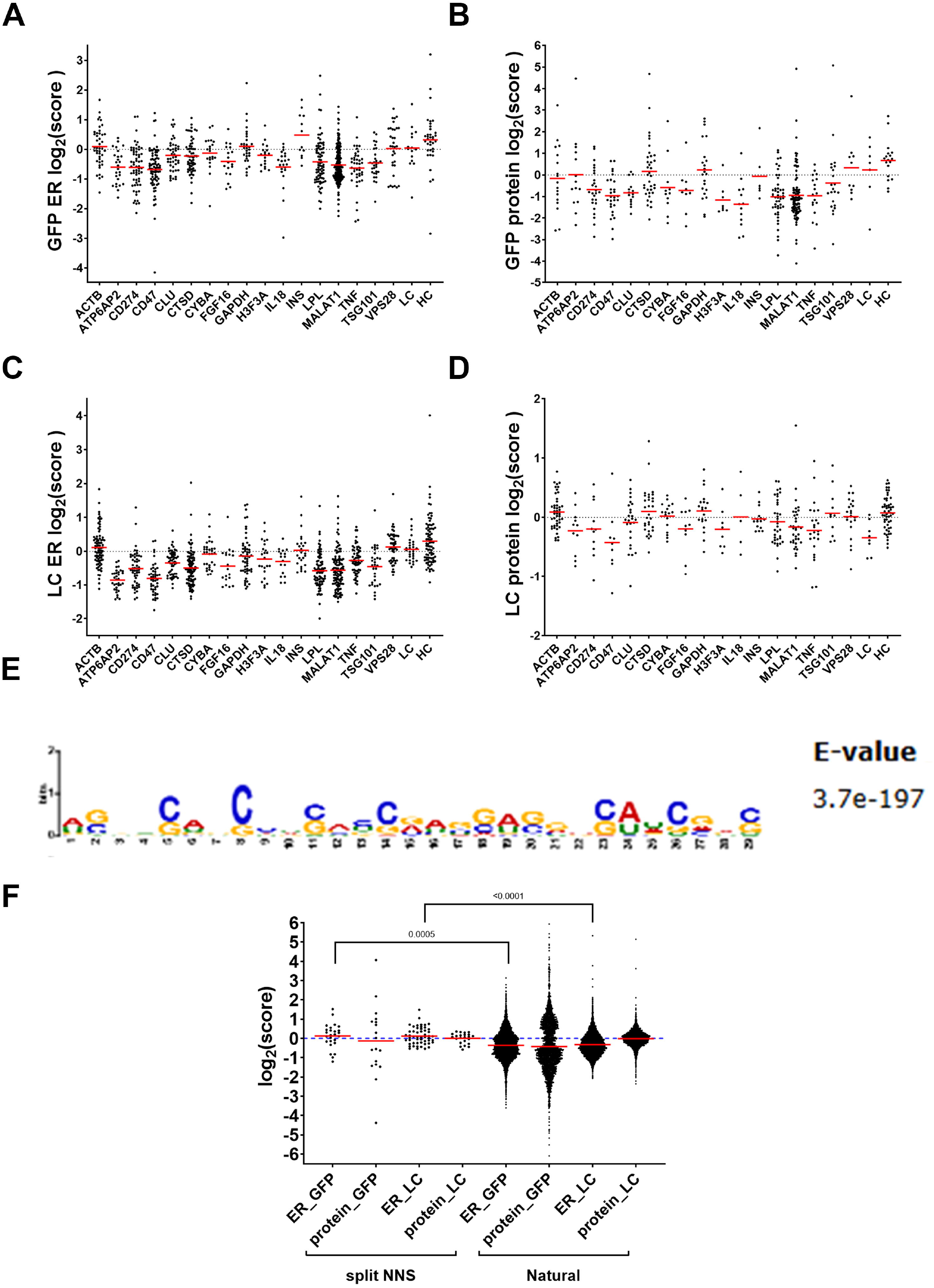
Score distribution of each gene tiling in the tiling sub-library and emerging split NNS motif. Graphs showing the score distribution of variants from each gene tiling sub-library. Each dot is a single variant. Note that IL2 was excluded due to very low number of variants (<5) with final scores. Red line is the average. See **File S10** for one-way ANOVA analysis **A)** Distribution of the GFP ER score. **B)** Distribution of the GFP protein score. **C)** Distribution of the LC ER score. **D)** Distribution of the LC protein score. **E)** MEME logo of the split NNS motif. **F)** Comparison of the scores of variants of the Natural SECReTE sub-library with split NNS (NNSx4-GAGGAG-NNSx3) to the entire Natural SECReTE sub-library (without the split NNS group). Significant p values are depicted (Welch’s t test, two-tailed; performed on the score, not log_2_(score)).

When comparing the two reporters, namely GFP and LC, for protein secretion scores (**Fig 7B and D**), we see a similar behavior of the tiling variants of most genes, with HC and VPS28 consistently enriched and most other secreted proteins depleted. INS protein scores were not enriched, whereas CTSD scores were enriched towards secretion. GAPDH scores were also enriched, whereas ACTB were slightly depleted with the GFP reporter, but not LC reporter. The LC protein scores also show enrichment in CYBA and TSG101 designs. However, one-way ANOVA analysis did not find any statistical significance between any of the tiling groups (**File S10**).

To gain further insight, we performed MEME analysis on the high scoring variants from the tiling sub-library. Out of 259 variants with any log_2_(score)≥0.5, we found 219 variants that contain a “split NNS” motif, *NNS*x4*-GAGGAG-NNS*x3, where *S* is *G* or *C* (**Fig 67E** and **File S11**). We then looked for variants from the Natural SECReTE sub-library that carry this motif. We found 72 variants that carry this motif and have a final score in at least one of the reporters and assays (**File S9**). We plotted the scores of the split *NNS* group and compared them to the sub-library of the Natural SECReTE (without split *NNS* containing variants) (**Fig 7F**). We found that split NNS variants show a mild, but statistically significant, enrichment in the GFP and LC ER scores, but not in the protein scores.

## Discussion

We developed a massively parallel screen to identify RNA motifs that affect mRNA partitioning between cytosol and membrane fraction, as well as the extracellular secretion of the encoded protein product (**Fig 1**). Using this system, we screened a designed library of 47,975 short sequences inserted at the 3’UTR of two GPI-anchored reporter proteins, GFP and IgG light chain (LC). This library was composed of natural and synthetic SECReTE motifs, and of various types of non-SECReTE sequences. Bioinformatic analysis of the two libraries and the two assays yielded results about the behavior of the different types of sequences in the library. In particular, we found that *RRY* repeats appear to confer higher scores for both reporters and both assays (**Fig 6**). Analysis of the library in the two reporters and in the two assays revealed two intriguing correlations. First, variants that confer high ER RNA localization tended to confer high protein secretion for the GFP reporter (**Fig 5A and C**). This result supports the basic notion that mRNA localization to the ER enhances protein secretion. However, the poor correlation for the LC reporter suggests that this phenomenon could be gene-dependent. It is possible and, perhaps, likely that the SECReTE element in the coding sequence of LC overcomes any potential additive effect of the 3’UTR library on protein secretion. In addition, we show in this study that endogenous mRNAs that encode secretome proteins from CHO cells are enriched in the ER membrane fraction and with SECReTE sequences (**Fig 2**), as previously published for other organisms (2, 19, 20, 44, 45).

We also found a positive correlation between the ER scores of both the GFP and LC reporters using both enrichment analysis (**Fig S8A, left**) and on a continuous scale (**Fig S8B**). This raises a hypothesis that some SECReTE variants might act similarly on different genes. The verification of this hypothesis will necessitate the use of additional protein reporters with this library. We also found a positive correlation in the protein score using enrichment analysis (**Fig S8A, right**). However, on a continuous scale the correlation decreases when comparing the protein secretion scores of the two reporters (**Fig S8C**). This low correlation could be either because of the internal SECReTE of LC or because of other differences in the reporter translation and/or protein properties. It could also be due to differences in the sample collection, since for the GFP we collected four categories (based upon GFP expression level and surface expression) whereas for the LC reporter we relied only on surface expression. Furthermore, we used only the high expression/high secretion fraction (**Fig S6**, p8) for the GFP protein scores. Thus, we are missing variants with low expression but showed high secretion (**Fig S6**, p6), which might affect the correlation between the two reporters.

Our analysis found an enrichment of *RRY* triplet repeats for high ER localization and protein scores as compared to the *NNY* repeat sub-library and all other sub-libraries (**Fig 6**). The *RRY* sub-library contains sequences with 39 repeats. However, even shorter *RRY* repeats could increase the enrichment in both reporters and both assays (**Fig 6**, “other RRY”). In addition to the RRY motif, we discovered another possible motif, which we called “split NNS”, with a mild but statistically significant positive effect on ER localization, but not on protein secretion (**Fig 7E-F**). This motif was discovered using our tiling sub-library, which offered an unbiased approach to find novel motifs. The tiling sub-library contained variants which tile the entire mRNA sequences encoding several proteins of choice, including secreted and non-secreted proteins. The depletion of CD247, CD47 and TNF variants was surprising, since it was previously shown that the mRNAs of these genes contain multiple A/U rich elements (ARE) that drive ER localization (46). In fact, most of the variants from secreted proteins were depleted (**Fig 7**). This does not preclude the idea that all the mRNAs examined are not preferentially localized to the ER, as opposed to having a cytoplasmic preference for localization. The split NNS motif appears to reside in flanking or slightly overlapping regions of SECReTEs (**File S9**, split NNS sheet). However, it is not clear if these flanking regions are required to enhance the effect of SECReTE, or work independently of SECReTE. The fact that the split NNS motif was found in the tiling sub-library suggests it is independent from SECReTE, but this will need to be tested in future research.

We used the Tomtom tool (47) of the MEME suit to look for motifs that are similar to the split NNS. The highest scoring (p = 0.000105) was *GGAGGA* that is similar to the middle part of the split NNS motif. The GGAGGA motif is recognized by the serine-arginine splicing factor 1 (SRSF1) protein. Although SRSF1 is mostly known for its splicing role, it is a nucleo-cytoplasmic shuttling protein that has a role in mRNA export (48) and in translation of a subset of mRNAs (49, 50). Thus, it is possible that SRSF1 binds the variants with a split NNS motif and promotes their nuclear export or translation on the ER.

Overall, the distributions of the variants from the various sub-libraries did not show a preference for SECReTE elements *per se*. For example, the polyR library and most of the gene tiling variants do not contain SECReTE sequences yet their variants seem to have similar distributions to SECReTE containing variants. Longer SECReTE motifs (*e.g*. double SECReTEs or the *NNY*/polyY sub libraries) do not seem to have an advantage over the Natural SECReTEs.

Several possibilities may account for these results. First, many variants were lost due to technical issues (see below) which reduced the statistical power but also may have lost a lot of useful information. Second, it is possible that SECReTE elements in the 3’UTR have a minor role or are non-functional when compared to SECReTE motifs present in the coding sequence. A recent publication suggests that about half of newly synthesized mRNAs transiently localize to the ER in a translation-independent manner, and that translation initiation inhibition increases this fraction (20). Thus, SECReTE motifs at the 3’UTR might be involved in promoting ER localization independently of the protein product, whereas 5’UTR and CDS SECReTE elements might maintain ER localization of secretome mRNAs. Third, both reporters have a SECReTE motif in their SP sequence, so any additional SECReTE may not necessarily increase mRNA ER localization or protein secretion levels. Hence, unless the added SECReTE is sufficiently robust in its action (*i.e.* with respect to mRNA localization to the ER or enhanced transcript stability), its effects could be insignificant. In addition, the fact that the LC reporter contains a second SECReTE sequence in the CDS might explain why the results with this reporter are less robust than those obtained with the GFP library. This may also explain any differences in sub-library scores associated with the top scoring variants of each reporter. Several other issues are listed in the *Limitations* section below. Other analyses we performed, such as looking for effect of GC/Y content, *NNY* length, positional source of the SECReTE element (*i.e*. 5’UTR, CDS or 3’UTR), mutations or flanking regions did not find any significant or consistent results worth reporting, probably due to subtle effects drowned in the background of the existing SECReTE or due to the other reasons mentioned above. However, the *RRY* and split *NNS* elements showed an effect strong enough to be detected in our assays. Yet these are restricted to the 3’UTR, and we do not yet know their effect when placed in the 5’UTR or CDS, or if they are superior to SECReTE at these positions.

While there may be >50,000 SECReTE variants contained within the protein-coding genes of the human genome (19), it is obvious that given their degenerative nature and limitations in the number of RBPs that many, if not most, motifs will be unrecognized. RBPs are expected to interact with the SECReTE zipcode in order to recruit the RNA to the ER. In yeast, for example, we found that ER-associated RBPs (*e.g.* Puf1, Khd1, Scp160, *etc*.) bind sets of mRNAs enriched with SECReTE motifs (19). Other work has shown that mammalian orthologs of these RBPs, for example Vigilin/HDLBP, also can bind sequences reminiscent of SECReTE (*e.g. CU* repeats) in secreted proteins and to enhance secretion (15). Given the plethora of SECReTE and perhaps other ER-motifs available in secretome proteins, we endeavored to identify *bona fide* functional versions of this motif using our assay. While some limitations remain (see below), nevertheless, this method represents a non-biased means for their identification, following which they could be used to help trap and purify SECReTE RBPs for further characterization and a better understanding of how RBP-mRNA interactions facilitate secretion at the level of the ER.

### Limitations

We suggest the following points for further improvement and refinement of future screens/experiments.

#### Toxicity

We found that both the reporter gene plasmids (*i.e.* with and without the oligonucleotide libraries) had some toxicity to the *E. coli* bacteria and to CHO cells, perhaps for different reasons. The toxicity in bacteria led to partial coverage of the library in cloning, an uneven distribution in the numbers of given variants, and to low plasmid yield. Thus, additional work should be invested in developing a less toxic reporter vector. We managed to reduce toxicity by flipping the whole expression cassette to its reverse orientation. In addition, the use of STBL-3 cells and their growth at lower temperatures decreased but did not abolish toxicity. This technical issue significantly hindered our ability to properly screen the libraries.

In CHO cells, GFP-GPI expression levels were significantly lower when compared to those of cytoGFP, without the SP and GPI anchor. Moreover, GFP-GPI-positive cells lost GFP expression rapidly over time. We suspect that the toxicity of the reporters in CHO cells stems either from the signal peptide (SP) and/or the GPI anchor used, since the cytoplasmic GFP vector showed very high expression and the cells did not show diminished GFP expression over time. Thus, it might prove beneficial to test other SPs or GPIs. Alternatively, it might be that the strong constitutive promoter caused an ER/Golgi stress due to high expression of a secreted/GPI-anchored protein. Thus, only cells with reduced expression survived. Such stress might have also affected ER-associated RBP recognition of SECReTE, which could lessen RNA association with the ER and thereby alter the differences between SECReTE variants. Changing to a weaker or perhaps an inducible promoter, one that might be induced just shortly before the assay is performed, might prove highly advantageous.

#### Reporter distribution

Cells expressing the reporter constructs alone showed wide cell-to-cell variability in expression (**Fig S6**). A large percentage of the population expressed the GFP reporter, although the GFP signal was not necessarily detected on the cell surface and GFP expression itself varied by at least two orders of magnitude. We suspect this is one manifestation of the reporter toxicity.

#### Cell fractionation

Cell fractionation showed that the reporter GFP-GPI mRNA was already highly associated with the membrane fraction (**Fig 3E**). Thus, it was probably difficult to detect major improvements in this regard. We do not know if the cause for this membrane association is due to the SP or to the SECReTE motif embedded in the SP coding sequence.

#### Cell surface detection

Microscopic imaging of the GFP reporter (**Fig 3A**) showed that it strongly decorates the cell surface. Thus, given this level of expression it might be difficult to detect increased secretion over this already high level.

#### LC reporter

With the LC reporter, we measured only one parameter: that of detection on the cell surface. In contrast, for the GFP reporter we could also detect the total level of expression, which was a useful parameter for judging overall expression. It might prove useful to include a fluorescent tag for the LC or, alternatively, develop a staining assay to detect total LC protein prior to sorting.

#### GPI anchor

A GPI anchor was used to attach the reporter proteins to the outer cell surface and in order to facilitate efficient sorting of the library. Yet, our assay was based on the assumption that SECReTE equally promotes the secretion of GPI-anchored proteins and other types of membrane proteins. Nevertheless, it is possible that a protein with a transmembrane domain or perhaps a soluble secreted protein would benefit more from an added SECReTE sequence, as we previously showed to be true for the latter in yeast (19).

#### 3’UTR

Our library was inserted in a fixed position at the 3’UTR. It is not clear if this has any advantage. It is possible that other positions or other 3’UTR sequences might prove more beneficial for specific SECReTE sequences.

#### Additional elements in flanking regions

Our screen focused on the SECReTE element. Although we did test the flanking region, some of our findings suggest that elements in the flanking region, or even sequences overlapping or within the SECReTE that are unrelated to the concept of SECReTE (e.g. split NNS) may have an additional effect on ER localization or protein secretion.

## Supporting information

all supplementary Files

## Acknowledgments

This work was funded by Lonza Biologics Ltd, who also supplied the IgG light and heavy chain plasmids and the CHO cells. We thank Ronnie Blecher and Hadas Keren-Shaul from the Nancy and Stephen Grand Israel National Center for Personalized Medicine (G-INCPM) at the Weizmann Institute of Science for their help with MARS-seq.

## Author contribution

G.H. constructed and tested the reporters, cloned the LC library, developed the assays and performed the experiments. G.L, S-T.Y, O.D., G.H. and Y.P. performed the bioinformatics analysis of the libraries. D.S. and T.O. designed the SECReTE library. D.S. sequenced and analyzed the plasmid library distribution. T.O. annotated the Chinese hamster 3’UTRs and analyzed the endogenous CHO mRNA distribution. F.G. and A.G. contributed to discussion of the results. J.E.G and Y.P. conceived the project and secured funding. G.H., G.L, S-T.Y., D.S., O.D. T.O., Y.P. and J.E.G. contributed to the writing of the paper.

## Supplementary Figures Legends

**Fig S1.**
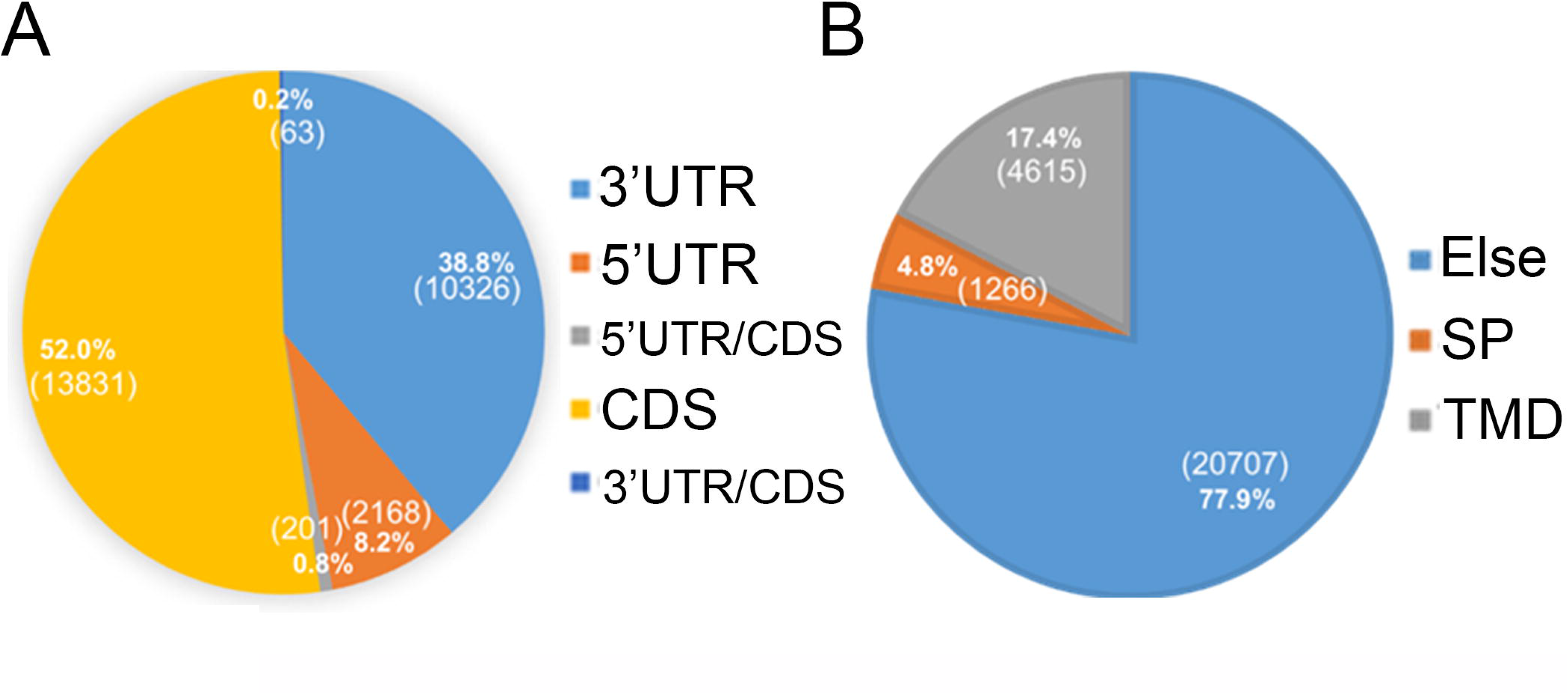
Distribution of natural SECReTE motifs along the mRNAs by physical or functional position. **A)** Pie chart depicting the distribution of SECReTE elements along the mRNAs by physical position. Numbers in parenthesis – number of motifs. **B)** Pie chart depicting the distribution of SECReTE elements along the mRNAs by functional position. Numbers in parenthesis – number of motifs. SignalPep – signal peptide; TM – trans membrane domain. Else – elsewhere along the mRNA.

**Fig S2.**
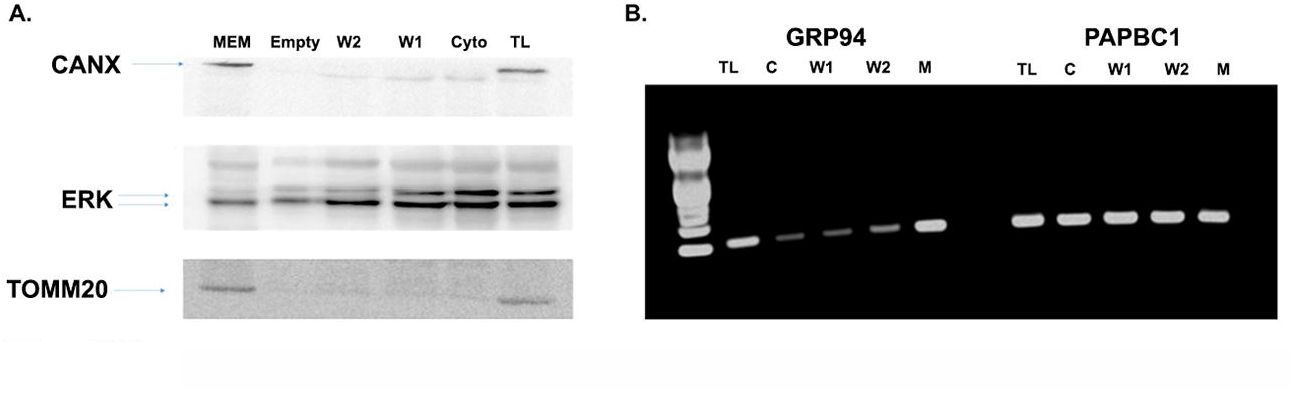
Distribution of cellular protein markers and mRNA upon cell fractionation. **A)** A Western blot showing the distribution of an ER protein (calnexin; CANX), a cytoplasmic protein (ERK), and a mitochondrial protein (TOMM20) upon subcellular fractionation. ‘cyto’ – cytoplasmic fraction ‘mem’ – membrane fraction. W1 and W2 are the supernatants of the two wash steps before membrane extraction. Empty – empty lane, showing leftover due to spill over of the sample from Wash2 well. **B)** An agarose gel (2%) showing the distribution across fractions of GRP94 and PABPC1 mRNAs by RT-PCR is shown. TL – total lysate; C-cytoplasm; M – membrane; w1, w2 – washes 1 and 2.

**Fig S3.**
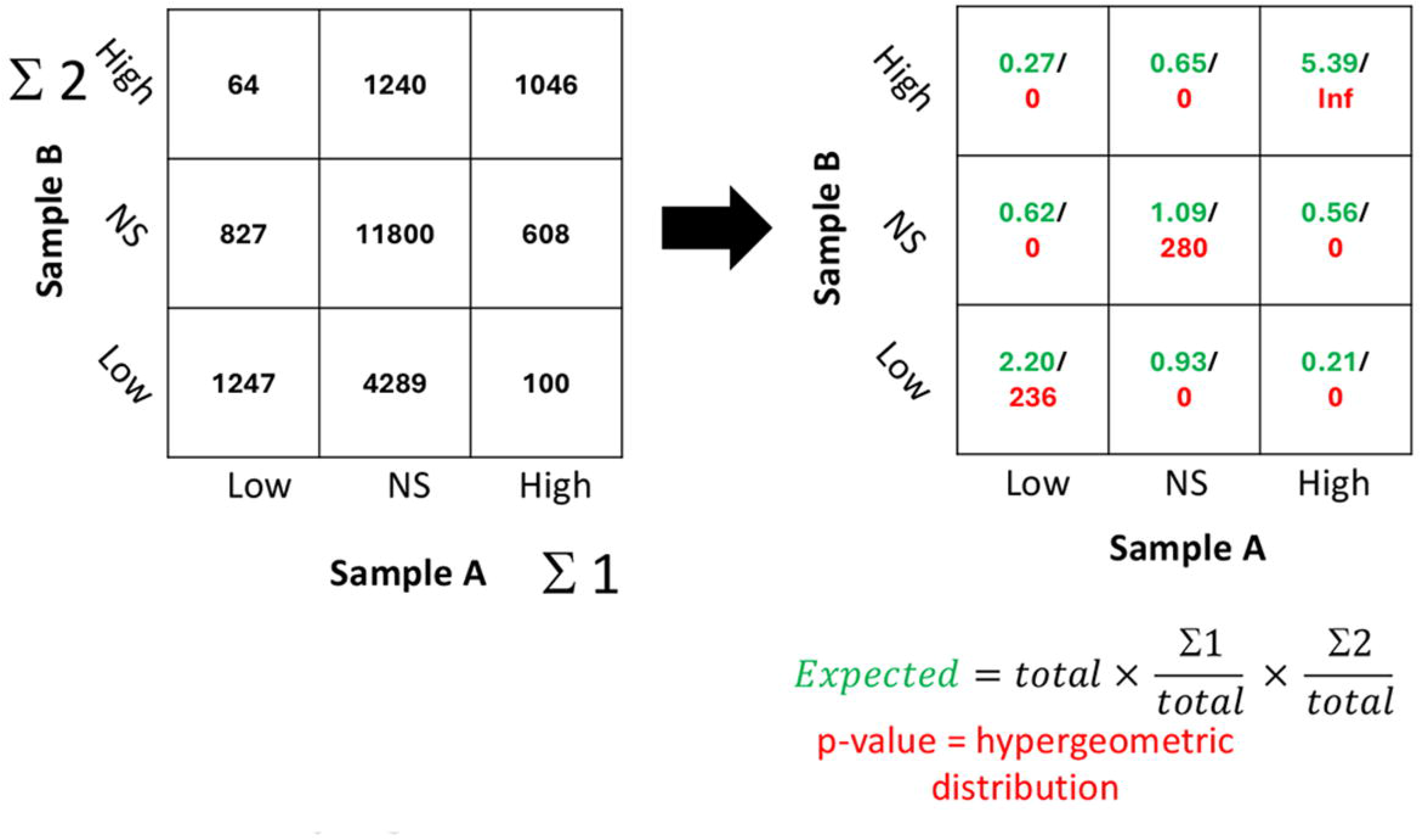
Mode of calculation of matrices of correlation between pairs of samples. On the left is a matrix of read counts of two hypothetical samples, A and B. The variants in each sample were classified into Low, High, and those that are not significant on either high or low (“NS”). The numbers in each matrix entry indicate how many library variants were on each joint class in the two samples (for example there were 1046 variants that scored highly on both samples, and 100 that scored highly on Sample A and lowly on Sample B). On the right are modes of calculation of over-representation and its hyper-geometric p value. In total from 21,221 designs. 1,754 variants scored High in Sample A and 2,350 on Sample B. Thus, ∼0.083 of variants scored high in Sample A and ∼0.111 in Sample B. Under the null hypothesis on no interaction, or correlation, between the two samples we would have expected that 0.083×0.111 of the 21,221 variants would have scored highly in both. This amounts to 194 variants expected. In reality, 1046 variants scored highly on both samples. This is 1046/194=∼5.4 folds higher extent of doubly high variants. The hypergeometric p-value of this overlap or higher is 10^-300^, shown in the matrix as -log 10 of p-value = 300. Hence in the upper right entry in the matrix on the right are the two values. Same calculation was done on every other entry in the right matrices.

**Fig S4.**
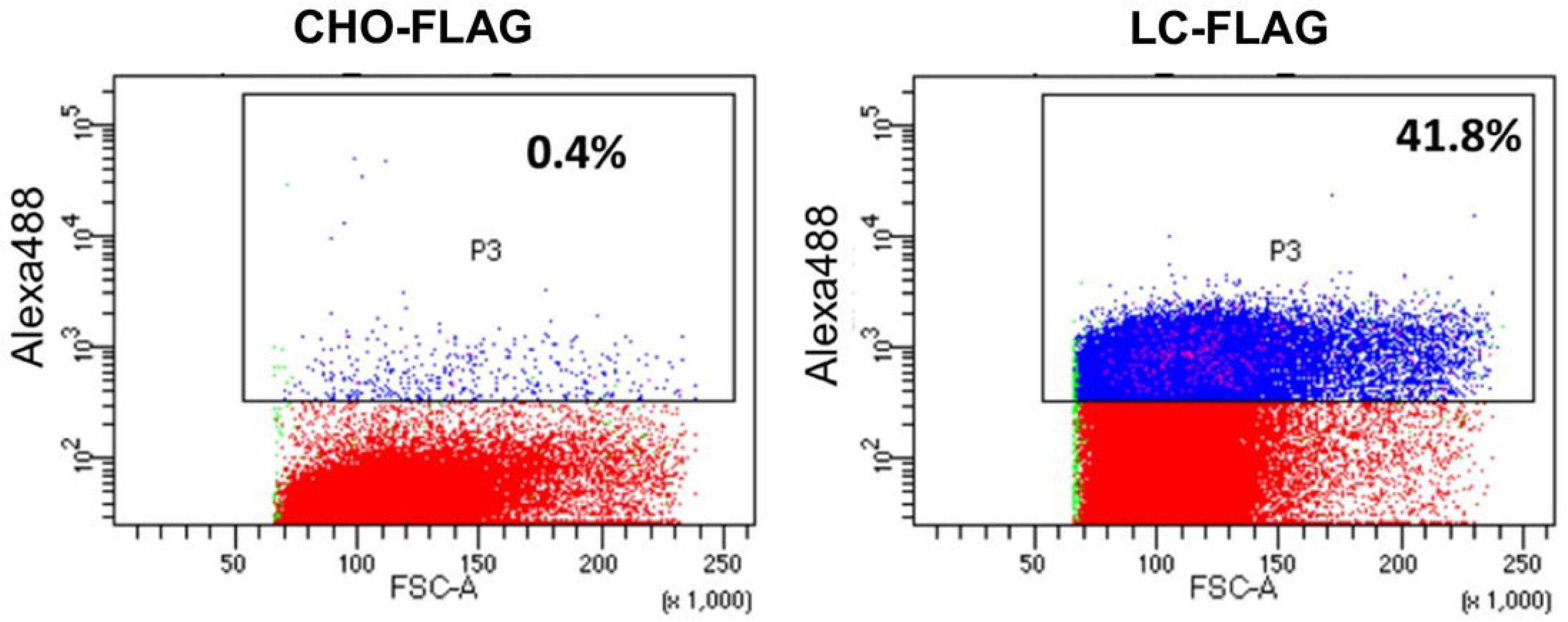
FACS analysis of CHO cells (left), or CHO cells transduced with LC-GPI reporter (right). The percentage of the P3 gate (high secretion) is indicated.

**Fig S5.**
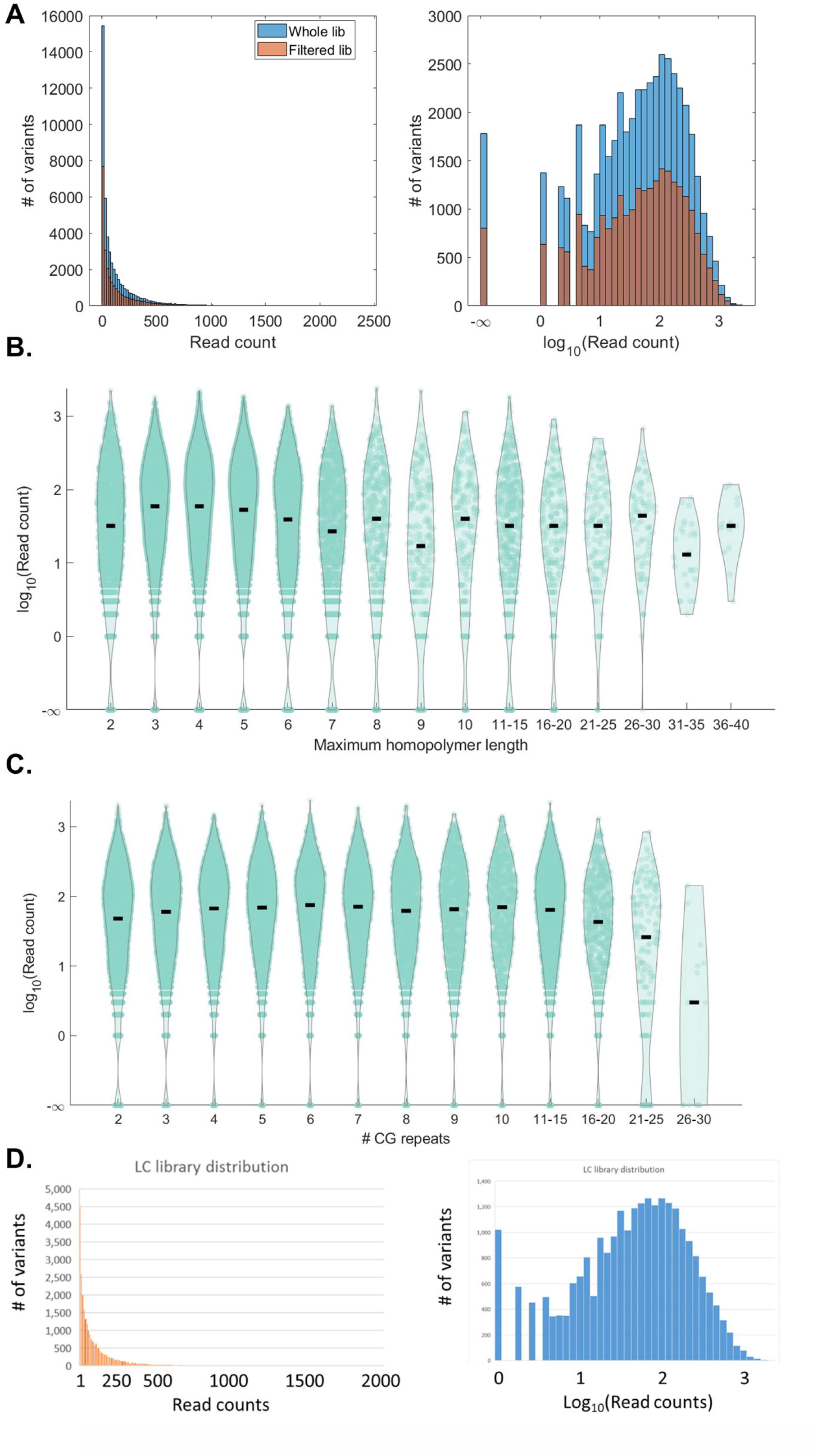
SECReTE variant distribution in the GFP and LC libraries. Sequencing of the GFP library plasmids shows the distribution of SECReTE variants in the library. **A)** The library distribution (blue) or the library after filtering variants with polyA/polyT>5 and CG repeats>6 (orange). Left – x-axis is read counts. Right – X-axis is log_10_(read counts). **B-C)** Read counts of variants versus nucleotide homopolymers **(B**) or CG repeats **(C)** length. **D)** Sequencing of the LC library plasmids shows the abundance of variants in the library. Left – X-axis is Read counts. Right – X-axis is log_10_(read counts).

**Fig S6.**
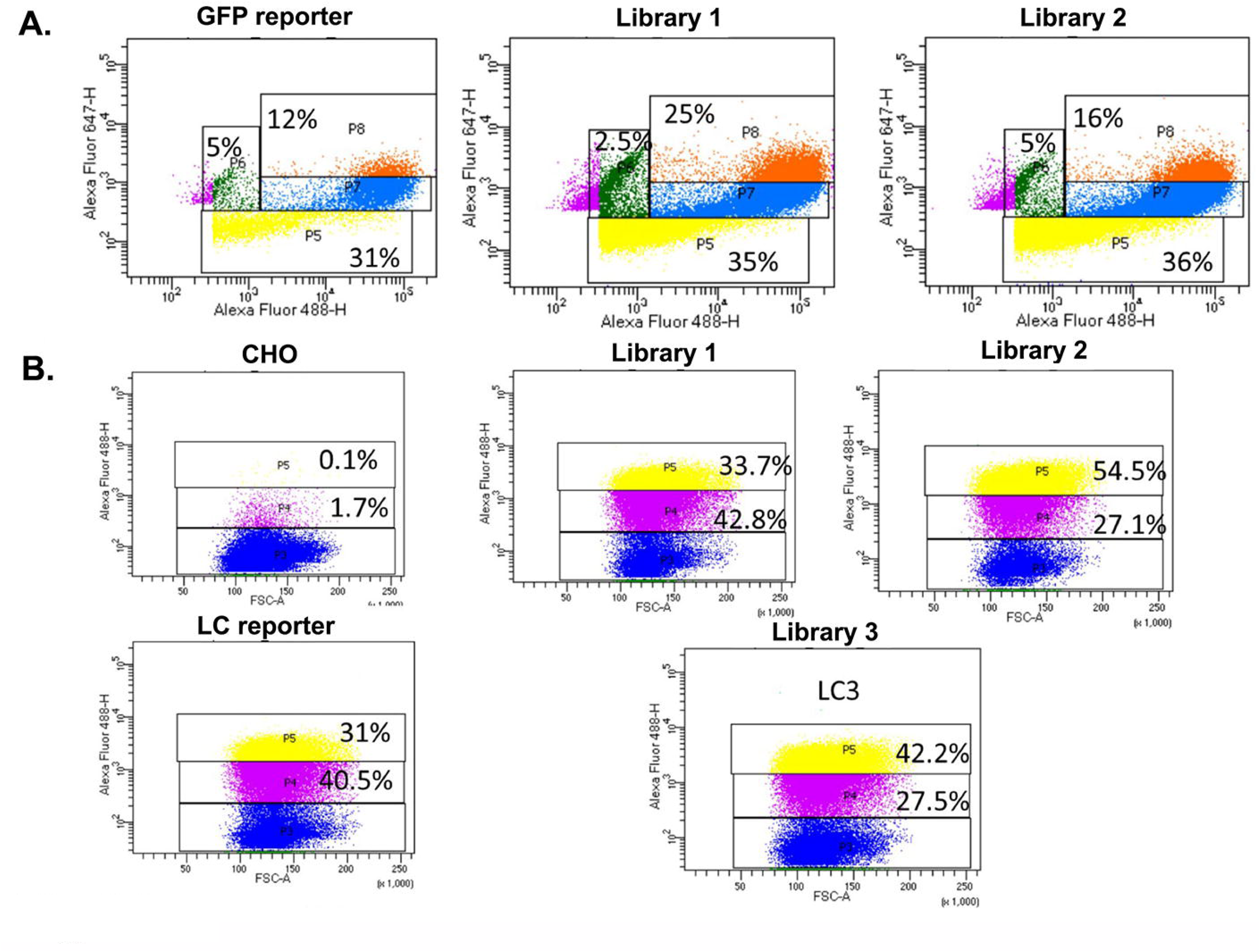
Gate selection for sorting of the GFP and LC libraries. Scatter plot analysis by FACS of CHO cells expressing the GFP **(A)** or the LC **(B)** reporter alone and the two new GFP libraries (GFP1, GFP2) and three LC libraries (LC1, LC2, LC3). For the GFP, X-axis depicts GFP fluorescence, Y-axis depicts A647 fluorescence. For the LC reporter, X-axis depicts the Forward scatter (FCS) and the Y-axis Alexa488 fluorescence (external FLAG-LC). CHO alone are shown for comparison to the LC library. The % of cells in the different gates are depicted.

**Fig S7.**
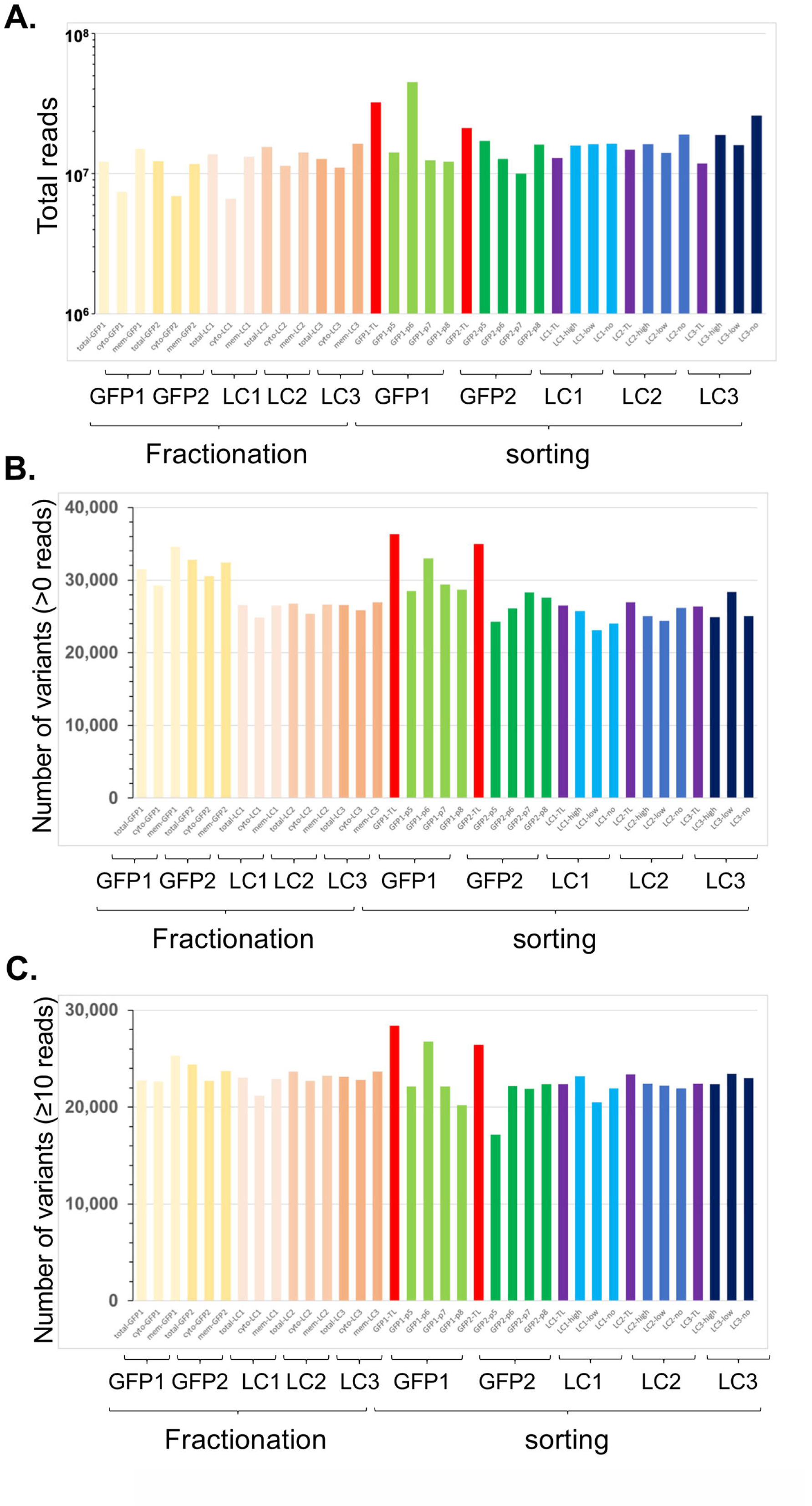
Sequence coverage of the experiments. Sequencing coverage of the two GFP reporter library and three LC reporter library biological replicates. (**A**) Total number of reads obtained for each sample for both the subcellular fractionation (Fractionation) and library sorting experiments (Sorting). (**B-C**). The number of variants with >0 reads (**B**) or ≥10 reads (**C**) that were detected. Red bars and purple bars – Total lysate of the GFP and LC libraries, respectively.

**Fig S8.**
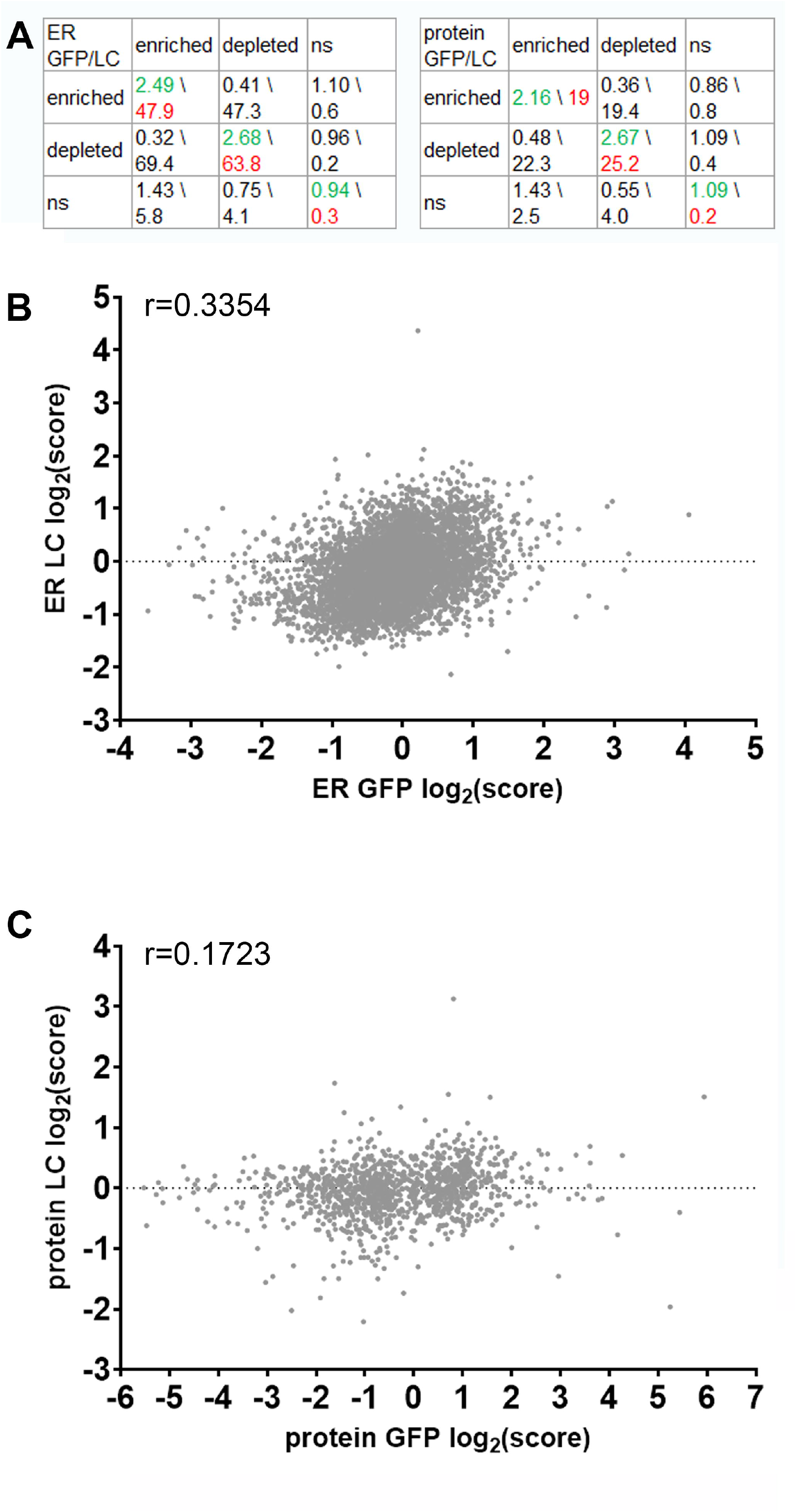
Positive correlation between GFP and LC reporters. (**A**) Enrichment analysis 3×3 tables of the ER scores (left) and protein scores (right). Each table compares the final scores of the GFP and LC reporter. Variants are classified into high (enriched), low (depleted), or non-significant (ns) changed levels in membrane-localized RNA or secreted protein levels, based on the relative presentations of variants (see Enrichment Analysis in Methods for details and **Fig S3** as an example). Numbers in green are the ratio of the observed to expected number of sequence variants overlapped between 2 categories for indicated samples. Numbers in red, -log10 of p-value of statistical significance for the overlaps from Hypergeometric test. **(B)** Graph showing GFP ER log_2_(score) (x-axis) *vs* LC ER log_2_(score) (y-axis). **(C)** Graph showing GFP protein log_2_(score) (x-axis) *vs* LC protein log_2_(score) (y-axis). Each dot is a single variant from the library. Pearson correlation (r) is noted at the top of the graphs.

**Fig S9.**
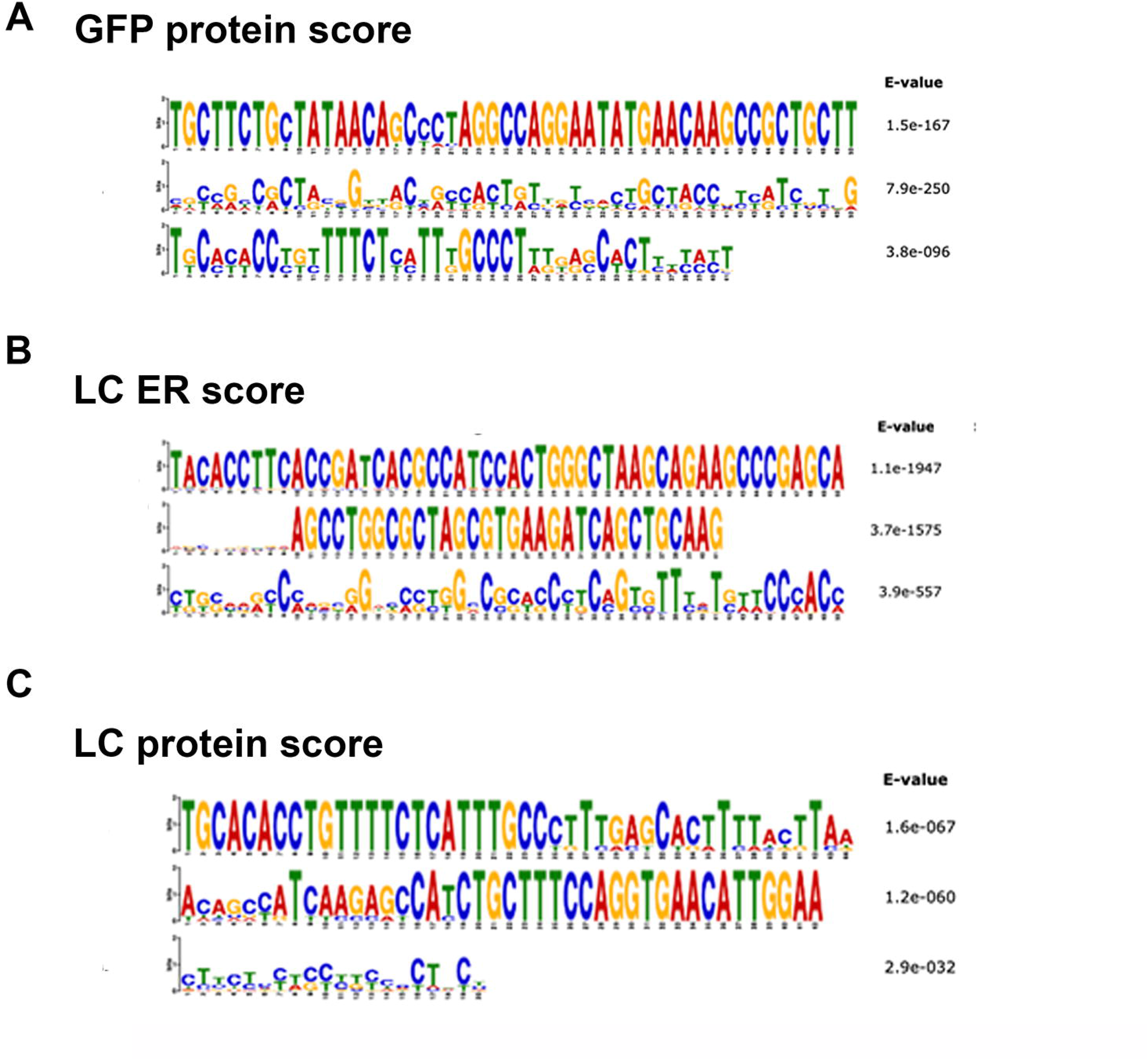
MEME derived RNA motif analysis of the GFP and LC reporter libraries. MEME motif finding algorithm was used on high scoring variants for each of the GFP reporter **(A)** or LC reporter **(B)** on each assay: ER RNA localization and protein secretion.

### Supplementary files

**File S1: list of human secretome genes and analysis of SECReTE elements of human secretome.** See Methods section for details.

**File S2: list of the SECReTE library.** See Methods section for details.

**File S3: The final gene model for Chinese hamster genome.** See Methods section for details.

**File S4: CHO mRNA MARS-seq data and distribution data for Fig 2**.

**File S5: sequencing reads of the GFP reporter library prepared by Twist Bioscience.**

**File S6: sequencing of the GFP reporter library after midiprep plasmid preparation in bacteria grown at 30°C or 37°C.**

**File S7**: sequencing reads for each variant in the GFP and LC libraries in each assay. cf = cell fractionation; cs = cell sorting; cyt - cytoplsmic fraction; mem - membrane fraction; TL - total cell; G1, G2 - GFP librarires 1 and 2; LC1, LC2, LC3 - LC libraries 1,2 and 3; p5, p6, p7, p8 - sorting gates of GFP libraries high, low - sorting gates of the LC libraries (see **Fig S7** for the sorting gates).

Note: the sum of each column is at the bottom of the column.

**File S8: ER and proteins scores of each variant in each replicate of the assays and reporters, and their average score.** 1 = no score. Average score in red font.

**File S9: The final scores of the variants, their log2(scores) and descriptive statistics of the sub-libraries.** See “readme” sheet for explanation.

**File S10: p-values for comparison of sub-libraries and genes of tiling library.** See “readme” sheet for explanation.

**File S11: a zip file of MEME output files in txt format.**

## References

1. Nyathi Y, Wilkinson BM, Pool MR. Co-translational targeting and translocation of proteins to the endoplasmic reticulum. Biochim Biophys Acta. 2013;1833(11):2392–402.

2. Cui XA, Palazzo AF. Localization of mRNAs to the endoplasmic reticulum. Wiley Interdiscip Rev RNA. 2014;5(4):481–92.

3. Lu J, Wu T, Zhang B, Liu S, Song W, Qiao J, et al. Types of nuclear localization signals and mechanisms of protein import into the nucleus. Cell Communication and Signaling. 2021;19(1):60.

4. Wiedemann N, Pfanner N. Mitochondrial Machineries for Protein Import and Assembly. Annual Review of Biochemistry. 2017;86(Volume 86, 2017):685–714.

5. Zilio E, Schlegel T, Zaninello M, Rugarli EI. The role of mitochondrial mRNA translation in cellular communication. Journal of Cell Science. 2025;138(9).

6. Baker A, Lanyon-Hogg T, Warriner SL. Peroxisome protein import: a complex journey. Biochem Soc Trans. 2016;44(3):783–9.

7. Irastortza-Olaziregi M, Amster-Choder O. RNA localization in prokaryotes: Where, when, how, and why. Wiley Interdiscip Rev RNA. 2021;12(2):e1615.

8. Eliscovich C, Buxbaum AR, Katz ZB, Singer RH. mRNA on the move: the road to its biological destiny. J Biol Chem. 2013;288(28):20361–8.

9. Buxbaum AR, Haimovich G, Singer RH. In the right place at the right time: visualizing and understanding mRNA localization. Nat Rev Mol Cell Biol. 2015;16(2):95–109.

10. Tian L, Chou HL, Fukuda M, Kumamaru T, Okita TW. mRNA Localization in Plant Cells. Plant Physiol. 2020;182(1):97–109.

11. Gonzalez I, Buonomo SB, Nasmyth K, von Ahsen U. ASH1 mRNA localization in yeast involves multiple secondary structural elements and Ash1 protein translation. Curr Biol. 1999;9(6):337–40.

12. Kislauskis EH, Zhu X, Singer RH. Sequences responsible for intracellular localization of beta-actin messenger RNA also affect cell phenotype. J Cell Biol. 1994;127(2):441–51.

13. Chou HL, Tian L, Washida H, Fukuda M, Kumamaru T, Okita TW. The rice storage protein mRNAs as a model system for RNA localization in higher plants. Plant Sci. 2019;284:203–11.

14. Mendonsa S, von Kügelgen N, Dantsuji S, Ron M, Breimann L, Baranovskii A, et al. Massively parallel identification of mRNA localization elements in primary cortical neurons. Nature Neuroscience. 2023;26(3):394–405.

15. Zinnall U, Milek M, Minia I, Vieira-Vieira CH, Müller S, Mastrobuoni G, et al. HDLBP binds ER-targeted mRNAs by multivalent interactions to promote protein synthesis of transmembrane and secreted proteins. Nature Communications. 2022;13(1):2727.

16. Koppers M, Özkan N, Nguyen HH, Jurriens D, McCaughey J, Nguyen DTM, et al. Axonal endoplasmic reticulum tubules control local translation via P180/RRBP1-mediated ribosome interactions. Developmental Cell. 2024;59(16):2053–68.e9.

17. Ma W, Mayr C. A Membraneless Organelle Associated with the Endoplasmic Reticulum Enables 3&#x2032;UTR-Mediated Protein-Protein Interactions. Cell. 2018;175(6):1492–506.e19.

18. Genz C, Fundakowski J, Hermesh O, Schmid M, Jansen RP. Association of the yeast RNA-binding protein She2p with the tubular endoplasmic reticulum depends on membrane curvature. J Biol Chem. 2013;288(45):32384–93.

19. Cohen-Zontag O, Baez C, Lim LQJ, Olender T, Schirman D, Dahary D, et al. A secretion-enhancing cis regulatory targeting element (SECReTE) involved in mRNA localization and protein synthesis. PLoS Genet. 2019;15(7):e1008248.

20. Child JR, Hofler AC, Chen Q, Yang BH, Kristofich J, Zheng T, et al. Examining SRP pathway function in mRNA localization to the endoplasmic reticulum. Rna. 2023;29(11):1703–24.

21. Kraut-Cohen J, Afanasieva E, Haim-Vilmovsky L, Slobodin B, Yosef I, Bibi E, et al. Translation- and SRP-independent mRNA targeting to the endoplasmic reticulum in the yeast Saccharomyces cerevisiae. Mol Biol Cell. 2013;24(19):3069–84.

22. Pyhtila B, Zheng T, Lager PJ, Keene JD, Reedy MC, Nicchitta CV. Signal sequence- and translation-independent mRNA localization to the endoplasmic reticulum. Rna. 2008;14(3):445–53.

23. Sarmah P, Shang W, Origi A, Licheva M, Kraft C, Ulbrich M, et al. mRNA targeting eliminates the need for the signal recognition particle during membrane protein insertion in bacteria. Cell Reports. 2023;42(3).

24. Jan CH, Williams CC, Weissman JS. Principles of ER cotranslational translocation revealed by proximity-specific ribosome profiling. Science. 2014;346(6210):1257521.

25. Costa EA, Subramanian K, Nunnari J, Weissman JS. Defining the physiological role of SRP in protein-targeting efficiency and specificity. Science. 2018;359(6376):689–92.

26. Thul PJ, Åkesson L, Wiking M, Mahdessian D, Geladaki A, Ait Blal H, et al. A subcellular map of the human proteome. Science. 2017;356(6340).

27. Consortium U. UniProt: the universal protein knowledgebase in 2021. Nucleic Acids Res. 2021;49(D1):D480–d9.

28. Safran M, Rosen N, Twik M, BarShir R, Stein TI, Dahary D, et al. The GeneCards Suite. In: Abugessaisa I, Kasukawa T, editors. Practical Guide to Life Science Databases. Singapore: Springer Nature Singapore; 2021. p. 27–56.

29. Stelzer G, Rosen N, Plaschkes I, Zimmerman S, Twik M, Fishilevich S, et al. The GeneCards Suite: From Gene Data Mining to Disease Genome Sequence Analyses. Curr Protoc Bioinformatics. 2016;54:1.30.1–1..3.

30. Whittle N, Adair J, Lloyd C, Jenkins L, Devine J, Schlom J, et al. Expression in COS cells of a mouse-human chimaeric B72.3 antibody. Protein Eng. 1987;1(6):499–505.

31. Broussau S, Jabbour N, Lachapelle G, Durocher Y, Tom R, Transfiguracion J, et al. Inducible packaging cells for large-scale production of lentiviral vectors in serum-free suspension culture. Mol Ther. 2008;16(3):500–7.

32. Jagannathan S, Nwosu C, Nicchitta CV. Analyzing mRNA localization to the endoplasmic reticulum via cell fractionation. Methods Mol Biol. 2011;714:301–21.

33. Kallehauge TB, Kol S, Rørdam Andersen M, Kroun Damgaard C, Lee GM, Faustrup Kildegaard H. Endoplasmic reticulum-directed recombinant mRNA displays subcellular localization equal to endogenous mRNA during transient expression in CHO cells. Biotechnol J. 2016;11(10):1362–7.

34. Schindelin J, Arganda-Carreras I, Frise E, Kaynig V, Longair M, Pietzsch T, et al. Fiji: an open-source platform for biological-image analysis. Nat Methods. 2012;9(7):676–82.

35. Jaitin DA, Kenigsberg E, Keren-Shaul H, Elefant N, Paul F, Zaretsky I, et al. Massively Parallel Single-Cell RNA-Seq for Marker-Free Decomposition of Tissues into Cell Types. Science. 2014;343(6172):776–9.

36. Trapnell C, Roberts A, Goff L, Pertea G, Kim D, Kelley DR, et al. Differential gene and transcript expression analysis of RNA-seq experiments with TopHat and Cufflinks. Nat Protoc. 2012;7(3):562–78.

37. Kohen R, Barlev J, Hornung G, Stelzer G, Feldmesser E, Kogan K, et al. UTAP: User-friendly Transcriptome Analysis Pipeline. BMC Bioinformatics. 2019;20(1):154.

38. Zhou Y, Zhou B, Pache L, Chang M, Khodabakhshi AH, Tanaseichuk O, et al. Metascape provides a biologist-oriented resource for the analysis of systems-level datasets. Nat Commun. 2019;10(1):1523.

39. Martin M. Cutadapt removes adapter sequences from high-throughput sequencing reads. 2011. 2011;17(1):3.

40. Zhang J, Kobert K, Flouri T, Stamatakis A. PEAR: a fast and accurate Illumina Paired-End reAd mergeR. Bioinformatics. 2014;30(5):614–20.

41. Langmead B, Salzberg SL. Fast gapped-read alignment with Bowtie 2. Nat Methods. 2012;9(4):357–9.

42. Danecek P, Bonfield JK, Liddle J, Marshall J, Ohan V, Pollard MO, et al. Twelve years of SAMtools and BCFtools. Gigascience. 2021;10(2).

43. Bailey TL, Johnson J, Grant CE, Noble WS. The MEME Suite. Nucleic Acids Res. 2015;43(W1):W39–49.

44. Reid DW, Nicchitta CV. Diversity and selectivity in mRNA translation on the endoplasmic reticulum. Nat Rev Mol Cell Biol. 2015;16(4):221–31.

45. Hoffman AM, Chen Q, Zheng T, Nicchitta CV. Heterogeneous translational landscape of the endoplasmic reticulum revealed by ribosome proximity labeling and transcriptome analysis. J Biol Chem. 2019;294(22):8942–58.

46. Ma W, Mayr C. A Membraneless Organelle Associated with the Endoplasmic Reticulum Enables 3’UTR-Mediated Protein-Protein Interactions. Cell. 2018;175(6):1492–506.e19.

47. Gupta S, Stamatoyannopoulos JA, Bailey TL, Noble WS. Quantifying similarity between motifs. Genome Biol. 2007;8(2):R24.

48. Müller-McNicoll M, Botti V, de Jesus Domingues AM, Brandl H, Schwich OD, Steiner MC, et al. SR proteins are NXF1 adaptors that link alternative RNA processing to mRNA export. Genes Dev. 2016;30(5):553–66.

49. Haward F, Maslon MM, Yeyati PL, Bellora N, Hansen JN, Aitken S, et al. Nucleo-cytoplasmic shuttling of splicing factor SRSF1 is required for development and cilia function. Elife. 2021;10.

50. Maslon MM, Heras SR, Bellora N, Eyras E, Cáceres JF. The translational landscape of the splicing factor SRSF1 and its role in mitosis. eLife. 2014;3:e02028.

